# Comparative assessment of gene drive release patterns and spread with a hex-based model for large-scale simulations

**DOI:** 10.64898/2026.07.04.736481

**Authors:** Jiahe Li, Chengwei Shi, Jackson Champer

## Abstract

Spatial population genetic and ecological modeling is often necessary to predict outcomes accurately. One example is gene drive, a rapid process involving spread of gene drive alleles through a population, usually to suppress pests or reduce transmission of vector-borne disease. Several existing models have been used to assess gene drive and other spatial processes. However, each of these has limitations, such as high computational cost and limited scalability, difficulty in incorporating environmental factors and complex lifecycles, or potentially simplified spatial structure. To overcome these challenges, we propose a hexagon-based computational framework that is designed to mimic continuous space for rapid genetic wave advances. This allows us to accurately simulate a larger spatial domain with lower computational investment. We implemented this model and compared the wave speeds of different gene drives with those obtained from other models. The results showed good agreement when hexagon width and dispersal were properly calibrated. We then determined optimal circular and linear (along roads) release patterns for a variety of gene drives and *Wolbachia* bacteria. To demonstrate the application of our framework to a hypothetical scenario, we constructed a model *Culex quinquefasciatus* mosquitoes on Hainan Island. We then evaluated the outcome of different gene drive release strategies, showing the transgenic insect release level necessary to achieve high gene drive coverage and how this could be further optimized based on mosquito and human distribution. Overall, our hex-based population genetic framework provides a flexible platform for realistic and large-scale models for gene drive and related applications.

## 1. Introduction

Gene drives are genetic elements that have the potential to bias their inheritance and spread through wild populations[1], [2], [3]. Engineered gene drives can be categorized into two main groups: suppression drives and modification drives. Suppression drives are utilized to decrease or eradicate populations of pest species, while modification drives are employed to spread beneficial traits throughout populations. Overall, gene drive systems offer a wide array of potential applications and hold significant promise in addressing public health and environmental challenges.

In recent years, significant advancements have been achieved[4]. Strong homing drives utilizing the CRISPR system have demonstrated effectiveness in both modification[5], [6], [7] and suppression[8], [9] efforts. Additionally, various confined toxin-antidote drives have been conceptualized[10], [11], [12], [13], [14], [15], with some showing promising results in *Drosophila* experiments[16], [17], [18], [19]. Further investigation is necessary to identify appropriate promoters and target sites, as well as to effectively engineer and test these novel drive systems in desired species. For deployment, it may be necessary to select the best design for a particular situation, with the level of confinement being an especially important consideration. This is usually assessed by computational modeling, which is needed to determine what might actually happen if a gene drive is released.

Gene drive systems can be modeled in panmictic populations as well as spatial arenas. Panmictic population models best represent situations where significant spatial structure is not present in the population. They can also apply to situations where spatial structure is not important to population dynamics, such as an even modification drive release over a spatially homogeneous arena. These models are also useful for gaining conceptual insights for new drive systems and identifying features such as density-dependent performance and release thresholds. Spatial models are likely more realistic in many situations, enabling simulation of spatially heterogeneous initial conditions, spatially specified releases, and changes in carrying capacity according to the landscape[20]. They can show additional properties of the drive such as wave of advance, as well as the outcome of a certain release pattern of the drive over time. This can be especially useful while testing proposed release scenarios and searching for the most efficient release pattern.

Among the different model frameworks, individual-based models show flexibility for accurately handling situations within continuous space[15], [21], [22], [23], [24]. This can allow study of complex phenomena such as chasing[22], [23], [25], [26]. However, individual-based simulations face limitations when scaling to larger and more complex populations. In these cases, their substantial computational demands make it difficult to simulate a large number of individuals in a spatially heterogeneous landscape. This can be partially ameliorated by different model architectures[27], but the complex genetics of gene drive will usually limit the study area to a relative small location, perhaps a few kilometers across.

Another method to incorporate spatial population structure is to divide a population into many subpopulations, each treated as a distinct deme[28], [29], [30], [31], [32], [33], [34], [35], [36]. Demes are linked by variable migration rates. This technique can be useful for modeling arbitrarily large areas, and it is often accurate when modeling separate islands linked by migration. However, certain critical spatial traits such as wave of advance are often not present in these models, even when they aim to represent larger, relatively continuous populations. This means that accuracy may be low in some situations, such as the rate of spread of gene drives.

Reaction-diffusion models represent another class of spatial gene drive simulations models[15], [37], [38], [39], [40], [41]. These models describe the change in the concentration of one or more substances distributed in space under the influence of local chemical reactions (which can represent reproduction and gene drive) and diffusion (which represents dispersal of individuals). While reaction-diffusion models offer powerful tools for understanding phenomena such as the formation of drive wave of advance and drive release dynamics, they often assume continuous space and deterministic processes. Therefore, it is hard to detect and analyze stochastic phenomena, such as chasing dynamics, with reaction-diffusion models. Additionally, solving reaction-diffusion models numerically still requires large computational resources if they need to examine small-scale processes when modeling large areas. Finally, such models can have difficulty in incorporating high heterogeneous natural landscapes.

In spatial modeling, square grids are often the preferred choice due to their ease of programming, straightforward segmentation, and extensive use in previous research. However, square grids are less effective in terms of connectivity because either only four directions of movement are permitted, or because the eight adjacent squares are not equidistant (diagonal squares are a factor of 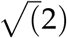 further away). This creates distortion in movement. In contrast, hexagonal grids have six directions of connectivity, meaning that each hexagon is surrounded by six equidistant hexagons with no other adjacent tiles. This reduces movement distortion and is particularly crucial for models involving migration between grids[42], [43], [44]. In image processing, hexagonal grids improve processing speeds by 25-50%[45], [46]. Compared to square grids, performance improved by over 40% improved with hexagonal edge detection operators[47]. Hexagonal grids are highly symmetrical and nearly circular, ensuring uniform connectivity among the six equidistant neighboring pixels. Conversely, square grids present a mix of edge-sharing and vertex-sharing neighbors, causing inconsistencies in boundary handling and reduced angular resolution in neighborhood pixels [47], [48], [49]. This geometric advantage advocates the adoption of isotropic hexagonal raster operators to avoid the anisotropy seen in square raster operators[50].

Considering the different model types, we aim to reduce computational costs while still maintaining crucial spatial properties by utilizing a hex-based model, wherein each hex is considered an independent deme but closely linked to its neighbors with high dispersal. The hexagonal grid structure allows for a more accurate representation of spatial dynamics, as it minimizes directional biases and better reflects natural movement patterns. This approach is similar to that used by existing tools such as HexSim[42], which has been successfully applied in ecological and population dynamics studies. It not only enhances model accuracy, but also enables us to simulate larger areas with realistic landscape and climate features due to having a lower computational burden, making it a potentially useful tool for studying gene drives in complex environments. We demonstrate the use of our model for optimal gene drive and *Wolbachia* release analysis with examples for *Culex quinquefasciatus* mosquitoes on Hainan Island.

## Methods

### A. Gene drive strategy

In this study, we investigate several gene drive systems, including homing drives, toxin-antidote drives, and CifAB drive. We also model *Wolbachia* with cytoplasmic incompatibility.

Homing drives utilize a nuclease, usually CRISPR/Cas9, to cut the wild-type allele in the germline of heterozygotes and induce homology-directed repair, which converts the wild-type allele to a drive allele[3], [51]. The probability that a drive/wild-type heterozygote converts a wild-type allele into a drive allele in the germline is the drive conversion rate. We consider resistance alleles formed by end-joining in both the germline (as an alternative to homology-directed repair) and the early embryo stages (from maternally deposited Cas9). The embryo resistance rate is defined as the probability that each wild-type allele in the offspring of a drive/wild-type heterozygote female undergoes transformation into a resistance allele. We assume that multiplexed gRNAs prevent formation of functional resistance alleles[52], [53], [54], [55], [56], [57], [58]. We model both homing modification and suppression homing drives. The modification drive targets an essential, haplosufficient gene, so nonfunctional resistance allele homozygotes are unable to survive. A recoded functional copy of the gene is provided in our drive allele, so individuals with drive alleles are always viable. The suppression drive targets a haplosufficient gene crucial for female fertility and does not have a recoded sequence. Thus, any females that lack a wild-type allele are sterile, including drive homozygotes.

We also study CRISPR toxin-antidote drives including Toxin-Antidote Recessive Embryo (TARE)[10], [16], [18], 2-locus TARE[11], Toxin-Antidote Dominant Embryo (TADE)[10], and TADE suppression[10], [11], [38], [59]. In all these systems, the drive is not copied by homology-directed repair, so all CRISPR cleavage forms disrupted/nonfunctional resistance alleles. TARE drives target an essential but haplosufficient gene, so disrupted allele homozygotes are nonviable. The drive has no threshold in ideal form but still has frequency-dependent performance. Cleavage happens in the germline, but we also model 100% embryo cleavage activity, where any alleles in a new offspring are cleaved if the mother has a drive allele due to maternally deposited Cas9 and gRNA. 2-locus TARE has two drive subtypes, each positioned at a separate locus. Each subtype provides rescue for the gene at their own locus, while targeting the wild-type allele at the other locus (each of these genes is also essential but haplosufficient). This produces an introduction threshold of 18% in the absence of fitness costs.

TADE targets and rescues a haplolethal gene, producing dominant-lethal disrupted alleles. One variant has cleavage restricted to the germline (with no introduction threshold, but frequency-dependent performance), and a more confined version has cleavage from maternal Cas9/gRNA deposition in the early embryo, producing an introduction threshold of 33% for ideal drive performance and no fitness costs. TADE suppression drives have additional gRNAs to target a female-fertility gene or are located inside the female fertility gene (disrupting it), while still providing rescue for the haplolethal target. We model this latter version, though both have the same population dynamics when performance is ideal[10]. This drive is also frequency-dependent with no introduction threshold for ideal drive performance, but it has distinct properties in spatial environments[38], [59].

Besides CRISPR-based drives, we examine *Wolbachia*, a bacteria capable of infecting arthropod species and being transmitted maternally. It can block pathogen transmission on its own[60]. Due to cytoplasmic incompatibility, when a male infected with *Wolbachia* mates with a female, viable offspring can be produced only if the female is also infected[61].The bacterium is transmitted maternally to all offspring and often carries a fitness cost. Most of our *Wolbachia* modeling is for an ideal system, but we model also in some simulations model the actual cost by setting the fitness of *Wolbachia*-infected individuals to 0.75, producing an introduction threshold of 31%[62]. We also examine a drive based on *CifA* and *CifB*, which is built from cytoplasmic incompatibility genes associated with a phage that infects *Wolbachia*, but integrated into the insect genome[15]. The CifAB drive system uses the same mechanism as *Wolbachia*. If a male drive carrier mates with a wild-type female, no progeny will be produced. The introduction threshold for CifAB drive is 37% for ideal performance[15].

Modification drives were released as homozygous individuals, and suppression drives were released as heterozygous individuals.

### B. Reaction-diffusion model

To compare our hex-based model to previous reaction-diffusion models of gene drive, we assess drive wave speed and shape using a 1-dimensional partial differential equation model. We run our simulations in an arena of length 300 with *dx* = 0.1, and *dt* = 0.001. In this model, we set carrying capacity *K* = 1 and define a generation as the time it takes for *K* individuals to be born and *K* individuals to die when the population is at carrying capacity. Denoting population size as *N* and the diffusion coefficient as *D, N*(*x, t*) is the total number of individuals at position x and time t. The population growth rate is calculated with the following equation, with low density growth rate *λ* = 5.

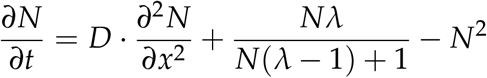

To avoid edge effects, we release drive homozygotes into the leftmost 20% of the arena and track the center of the resulting wave. The wave’s center is the point of 50% allele frequency (or 50% population suppression for suppression drives). We then calculate the wave’s speed by finding its stable velocity. In all of these tests, drive parameters are optimal, with the exception of the *Wolbachia* fitness cost.

### C. General hex model

In our hexagonal model, we construct an *m* × *n* grid of hexagons. Each hexagon is defined by its position and the number of individuals (though note that this is a continuous variable in our current implementation) of each genotype it contains. We implement a migration structure based on a diffusion model, with an adjustable average dispersal distance. Dispersal is modeled using a 51×51 diffusion kernel that approximates a normal distribution. We limit movement to a maximum of 25 hexes per weekly time step (extremely few individuals would be capable of moving beyond this distance based on our dispersion kernel). An example of our diffusion kernel with an average dispersal of 11.211 hexes (based on mosquito dispersal, see below) is shown in Figure 1.

**Figure 1.**
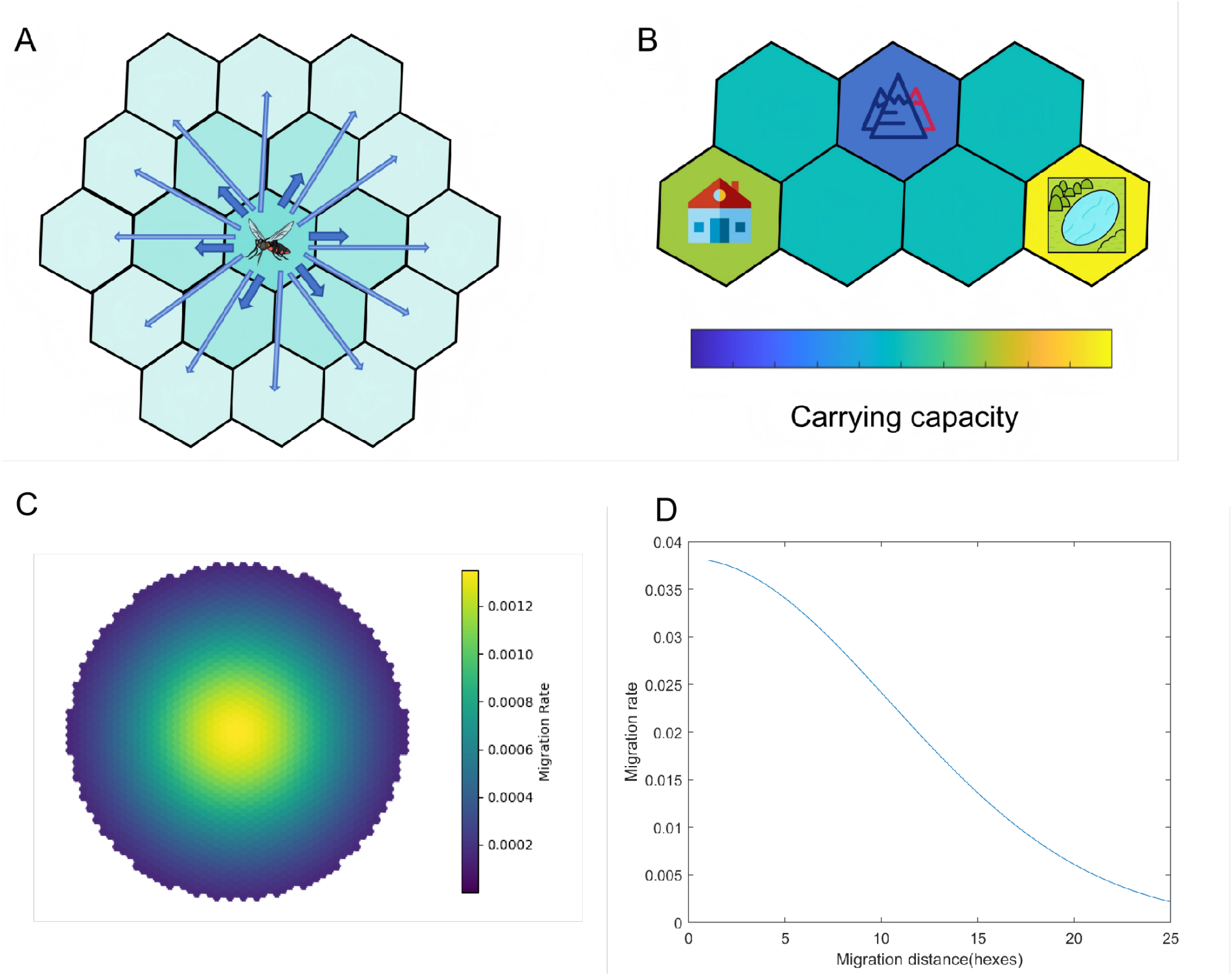
The hex model. (A) In each time step, adult mosquitoes can move to nearby hexes. Dispersal patterns are based on a normal distribution. (B) The carrying density of each individual hex is based on several factors, including human population density, water sources, temperature, and altitude. (C) The exact dispersal kernel for *Culex* mosquitoes. The migration rate refers to the proportion of mosquitoes from the central hex that migrate to the shown hex. (D) The dispersal kernel shows migration rate as a function of distance in hexes. 25 is the maximum migration distance in this model.

**Figure 2.**
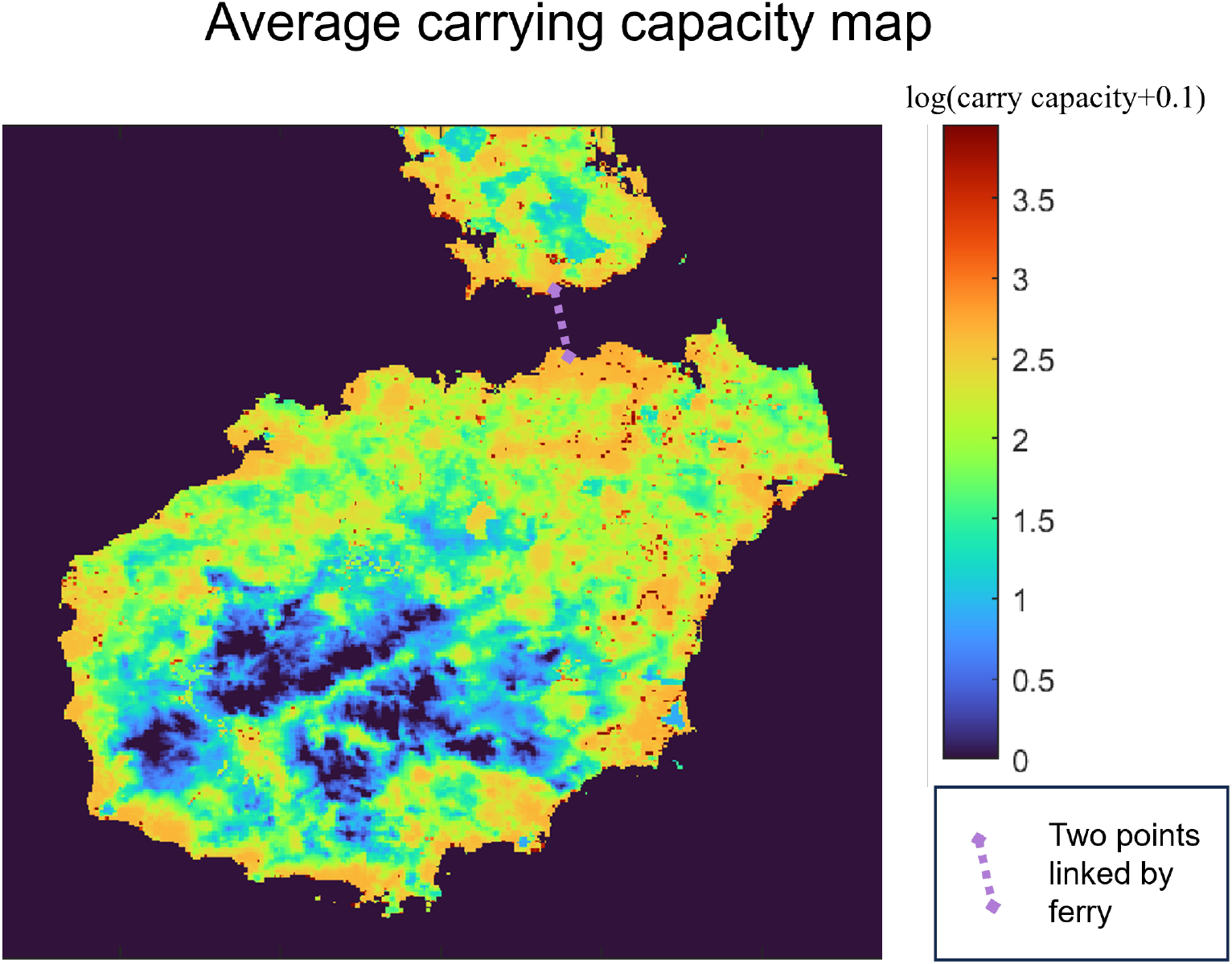
Mosquito carrying capacity of Hainan Island. Color shows the relative mosquito population density, averaged across all months. Migration of mosquitoes northward to the mainland is possible via ferry and is marked in purple.

Unlike reaction-diffusion models, in all of our hex models, time is explicitly discrete. In the general hex model, each time step represents one generation. The equations and parameters of reproduction are similar to the reaction-diffusion model, with the only difference being Δ*t* = 1 in most cases. We use our basic hex model for drive wave speed assessments and radial drop spread assessments.

For wave speed assessments, our arena size is 300 × 300. Initially, drive homozygotes cover the left 20% of the arena, and wild-type covers the rest. We set our checkpoints at 50% and 60% of the arena and calculate our wave speed like in the reaction-diffusion model, with optimal drive parameters unless otherwise specified. We also examine radial release scenarios, which are simulated in a 500 × 500 arena. Our radial release is characterized by *release radius* and *release f requency*. Initially, the whole arena is covered with wild-type with a population density of 1 in each hex, and we release drive homozygotes in each hex within a distance *release radius* from the center of the arena at the frequency of *release f requency*. We release drive homozygotes (for modification drives) or heterozygotes (for suppression drives) of both sexes. As time passes, the drive will advance outward radially, or in some cases contract for confined drives with small release sizes. In the first five generations, we set Δ*t* = 0.01 to prevent population sizes from becoming negative (the drive release initially creates an area of high density, increasing the death rate), and we keep Δ*t* = 1 for the rest of the simulation. In our two-year time points, which is 33 generations (see mosquito model subsection), we calculate the maximum radius in which the drive carrier frequency is at least 50% for modification drives, and the maximum radius in which wild-type frequency is below 50% for suppression drives.

### D. Mosquito hex model

To model mosquitoes, we extend our general hex model based on previous work for *Anopheles* [63], but reparameterized to consider *Culex* mosquitoes, which share a similar life cycle but have higher adult survival and dispersal distances[64].

Time steps in the model represent one week. We incorporate two juvenile stages (*juvenile*_0_, *juvenile*_1_), eight adult female stages (*female*_0_– *female*_7_), and five adult male stages (*male*_0_–*male*_4_), reflecting the longer lifespan of female mosquitoes.

Each week, a fraction of adults survive based on their age *i*. The survival rate for *female*_*i*_ is 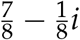 and for *male*_*i*_ is 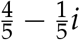. Survival rates for *female*_7_ and *male*_4_ are zero, indicating maximum lifespans of eight weeks for adult females and five weeks for males. In this model, adult mosquitoes do not experience density-dependent death rates.

At carrying capacity, each fertile female produces *eggs* = 40 per week, approximating the number of potentially viable eggs produced in the wild assuming that females must hunt for blood meals, find places to lay eggs, and have some offspring be nonviable even under low-density conditions due to predation or other factors. The survival rate of juveniles *juvenile*_*i*_ is denoted by *survival*_*i*_, with juveniles experiencing density-dependent competition:

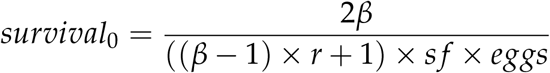

The intrinsic growth rate is *β* = 5.

To maintain populations at equilibrium with adult female carrying capacity *K*, we explicitly model density-dependent competition among age-one juveniles. The competition ratio *r* is defined as the ratio of actual juveniles to the expected number at carrying capacity:

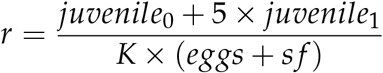

The term *s f* calibrates the model to ensure constant generation size:

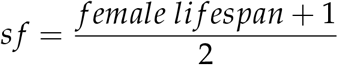

We assume *survival*_1_ is a constant set to 1 (thus modeling all larval mortality in the first week). Thus, new adults are produced as *juvenile*_1_ × *survival*_1_ and are equally divided into *female*_0_ and *male*_0_.

We perform drive wave speed and drive spread radius assessments for the mosquito hex model similarly to the general hex model. The key difference is that only age-one-adult drive individuals are released initially, and only adults are considered when calculating drive allele and carrier frequencies.

We examined cost-effective release strategies to maximize the number of drive alleles after a fixed time for each drive allele in the initial release. We define release efficiency as the ratio of covered area (with drive individuals) to released mosquitoes:

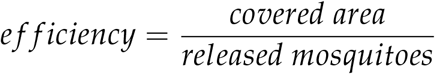

### E. Hex model of Hainan Island

*Culex pipiens* is the most prevalent mosquito species in Hainan and is capable of transmitting several severe diseases[65] such as West Nile Virus, Usutu Virus, filariasis, encephalitis, and Avian Malaria. We thus applied our *Culex pipiens* model to the Hainan region.

We utilized 2599 × 2601 hexagons to model Hainan Island and part of Guangdong Province. Each hexagon has a diameter of 109.55 meters, and the migration rates distribution is also the same as the mosquito model. We set the average dispersal distance equal to 1.1121*km*[66]. To model potential mosquito migration between Hainan and Guangdong via ferries, we incorporated special migration between two coastal points on Hainan and Guangdong, with a “ferry migration rate” of *mr*_*b*_ = 0.01.

To generate the carrying capacity of each hex, we employed a layered structure to account for various terrain factors[67], [68]. Our model is similar to a previous one used for *Anopheles gambiae* in Africa[69], but several aspects are modified to improve realism. Specifically, we used human population density instead of number of human settlements as our input parameter for determining mosquito capacity. We also included effects of temperature and altitude. High altitude and temperature extremes will negatively influence carrying capacity. Additionally, we considered separate factors as multiplicative for carrying capacity, so a severe problem in a single category would be sufficient to prevent significant numbers of mosquitoes from being present.

- Rainfall

The influence of rainfall, *rain f unc*, is a function of *rain f all*(*mm*). (*a*_1_, *p*_1_, *p*_2_ are parameters)

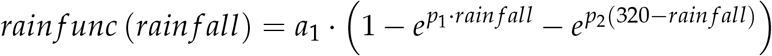

- Water

The influence of water, *water f unc*, is a function of permanent water (*water*_*p*_) and seasonal water (*water*_*s*_). If there exists water, then *water*_*p*_ or *water*_*s*_ equals 1, else 0. (*a*_2_, *p*_2_, *p*_3_ are parameters)

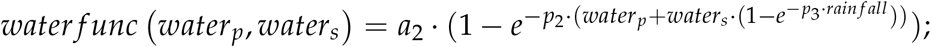

- Human density

The influence of human population, which makes it easier for mosquitoes to find blood meals, *bloodmeal*, is a function of human population density(*people*/*km*^2^), noted as *pop*. This function is set to prevent excessive numbers of humans from significantly influencing mosquito density, once number of humans is no longer limiting. (*a*_3_, *p*_4_, *p*_5_ are parameters)

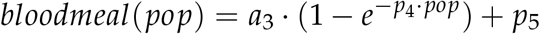

- Altitude

The influence of altitude, *altitude f unc*, is a function of *altitude*(*m*). Because our modeled mosquitoes cannot survive at altitudes higher than 800 meters, we set *altitude f unc*(800) to zero. (*a*_4_, *p*_6_ are parameters)

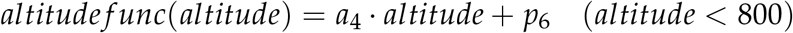

- Temperature

Mosquito viability is optimal with intermediate temperature, so we considered the monthly maximum and minimum temperatures. The influence of temperature(°C), *temp f unc*, is a function of temperature (*temp*), monthly maximum temperature (*t*_*max*_), and monthly minimum (temperature (*t*_*min*_). (*a*_5_, *p*_7_, *p*_8_, *p*_9_, *p*_10_, *p*_11_, *p*_12_ are parameters)

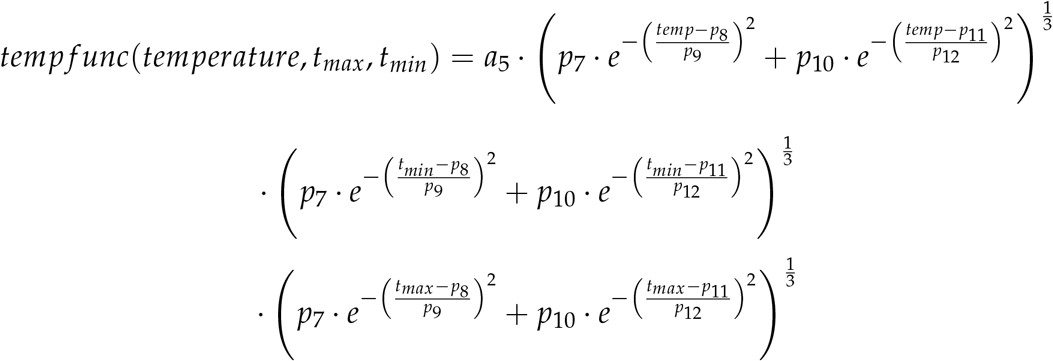

- Summary

Combining all the different factors, we obtain

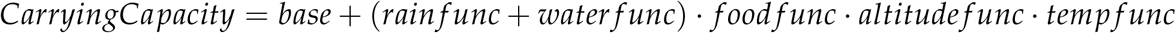

where *base* is the baseline mosquito population, set to 0 in our model but adjustable under different circumstances. In this model, the mosquito population has an average of 2.54 × 10^7^ in arbitrary units, with seasonal maxima and minima of 3.54 × 10^7^ and 8.05 × 10^6^, respectively. Each hex has an average of 7.6165 adult mosquito units.

### F. Data collection

We collected data using the High-Performance Computing Platform of the Center for Life Science at Peking University. Notably, many Hainan Island models could be completed in under two hours (though the more complex 2-locus TARE drive took longer). We used MATLAB R2022a to process data and prepare figures. All MATLAB models are available on GitHub (https://github.com/Haruka123123/Hex-model).

## 3. Results

### A. Wave of advance

One critical aspect of our hex-based model is that it can recapitulate drive wave formation (as seen in reaction-diffusion and individual-based continuous space models) despite being composed of multiple discrete demes. To assess this, we visualized the wave shape and checked the wave speeds for all of our different genetic systems. We adjusted the diffusion coefficient *D* to match the average dispersal of the hex model *avd*.

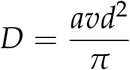

In the hex model, we analyzed drive wave formation in two different forms: one where the drive wave advances perpendicular to the flat sides of the hex, and one where it advances 60 degrees off from this direction toward the junction points of three hexes (Figure S3). In general, the hex model (both directions) and PDE model were very closely matched in wave speed if the average dispersal level was at least one hex per generation (Figure 3). However, for shorter range dispersal, the hex model starts to diverge from the PDE model, yielding higher wave speeds that eventually become lower and then zero for threshold-based drive systems. This is the critical point where the system loses the ability to mimic wave formation in continuous space. For the ideal homing and TARE drives, which have zero introduction threshold, the drive could nonetheless continue to advance with arbitrarily low dispersion. However, underdominance systems with higher thresholds, including TADE and CifAB, exhibited a critical dispersal threshold below which the wave failed to advance. This shows character of linked deme systems rather than continuous space. We can thus conclude that with sufficiently high dispersal per time step, the hex model will have close wave speeds to continuous space models. One important exception is homing drives, where wave advance is driven by the drive at low frequency (“pulled waves”). For this system, discrete time steps tend to slow the wave down, preventing convergence for higher dispersal. However, all discrete-time models are affected by this phenomenon compared to continuous time models such as our reaction-diffusion model, and indeed, reality may also often diverge from the continuous reproduction and movement assumptions of continuous time models. It remains unclear which may better approximate a real-world species for a certain situation, pending testing or availability of detailed ecological information.

**Figure 3.**
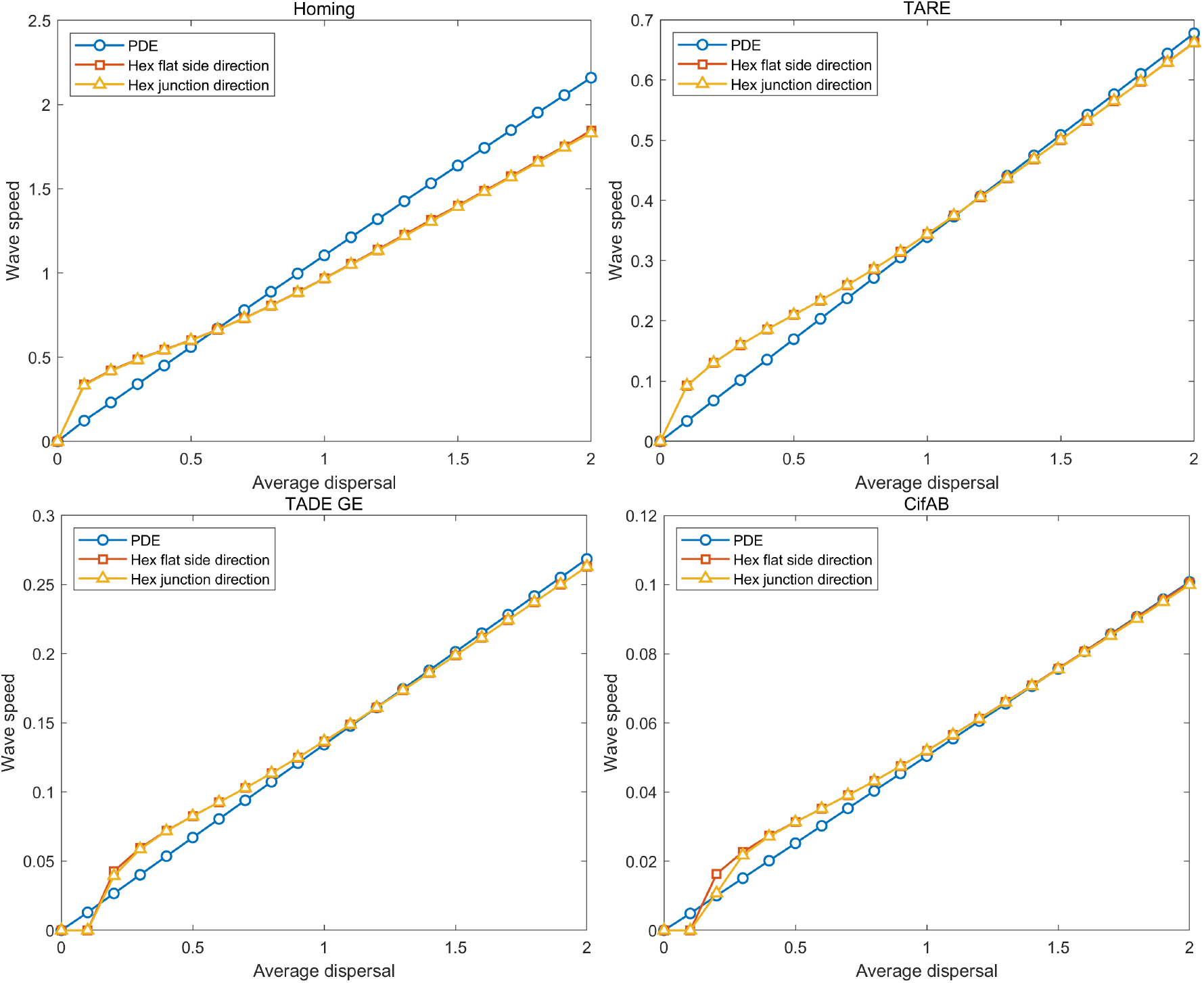
Wave speed comparison. For homing modification drive, TARE drive, TADE GE drive, and CifAB drive, we show the drive wave speed for two directions in the discrete-generation hex model and in the reaction-diffusion Partial Differential Equation (PDE) model. Wave speed is a function of the average dispersal distance.

In addition to the wave speed, we can use wave shape (drive frequency as a function of distance) as a potential point of comparison between the hex model and a reaction-diffusion model, to see what level of distortion can be expected in the hex approximation. For the same four drives, we found that at low dispersal (0.5 hexes per generation) wave shapes are fairly close, but still somewhat distorted, tending toward more intense drive gradients in the hex model (Figure S5). For high dispersal (10 hexes per generation), the wave shapes were nearly identical between the different models, with only TADE drive with embryo activity having a notable difference (in this case, the gradient was slightly less intense in the hex model).

To gain an understanding of different drives’ performance in our hex model, we adjusted parameters of our drives to model imperfect but realistic scenarios. These variable parameters were somewhat drive-dependent and included drive conversion, germline and embryo cut rates, fitness costs, and dispersal. We used these simulations to collect the drive wave speed (Figures S6-7). In some cases involving high threshold drives and high fitness cost, the wave of advance was unable to form. In most cases, though, we successfully measured wave advance speeds, which were similar to previous results using an individual-based model[21] and within a rounding error of the reaction-diffusion model with relatively high dispersal value, aside from the slightly higher wave speeds for homing drives in the reaction-diffusion model.

### B. Optimal circle drive release

To examine optimal drive release strategies using our hex model, we examined circular releases varying in three parameters: *Release Duration, Relative Release Frequency*, and *Release Radius*. We released gene drive/*Wolbachia* individuals beginning in the first generation and continuing for the period defined by the *Release Duration*. The amount of each release is determined by the *Relative Release Frequency*, which is a set ratio compared to the local wild population size in each release hex. All releases were conducted within a circular area defined by the *Release Radius*. We define efficiency as the coverage area divided by the release number. Coverage Area usually represents the total area of all hexagonal zones where more than 90% of the population carries the gene drive. While some drive alleles will spread beyond this area, we elected to use 90% to represent the area with strong drive protection. Initial data for drive spread examined the effect of varying release effort (Figures S8-9).

The efficiency calculation was performed at different time points depending on the type of gene drive used. For homing drives (including homing modification and homing suppression), where drive spread is rapid even with small releases, efficiency was calculated at the end of year 2. For all other drive types, which have substantially larger releases and slower spread, efficiency was calculated at the end of year 5. The spread radius for each drive is presented in Table 1. Note that we exclude CifAB and *Wolbachia* with fitness costs from this sort of analysis because they take a long time to form a wave and have a very slow speed. Though effective at modifying populations where they are released, their optimal release will nearly always be over the entire arena.

**Table 1.**
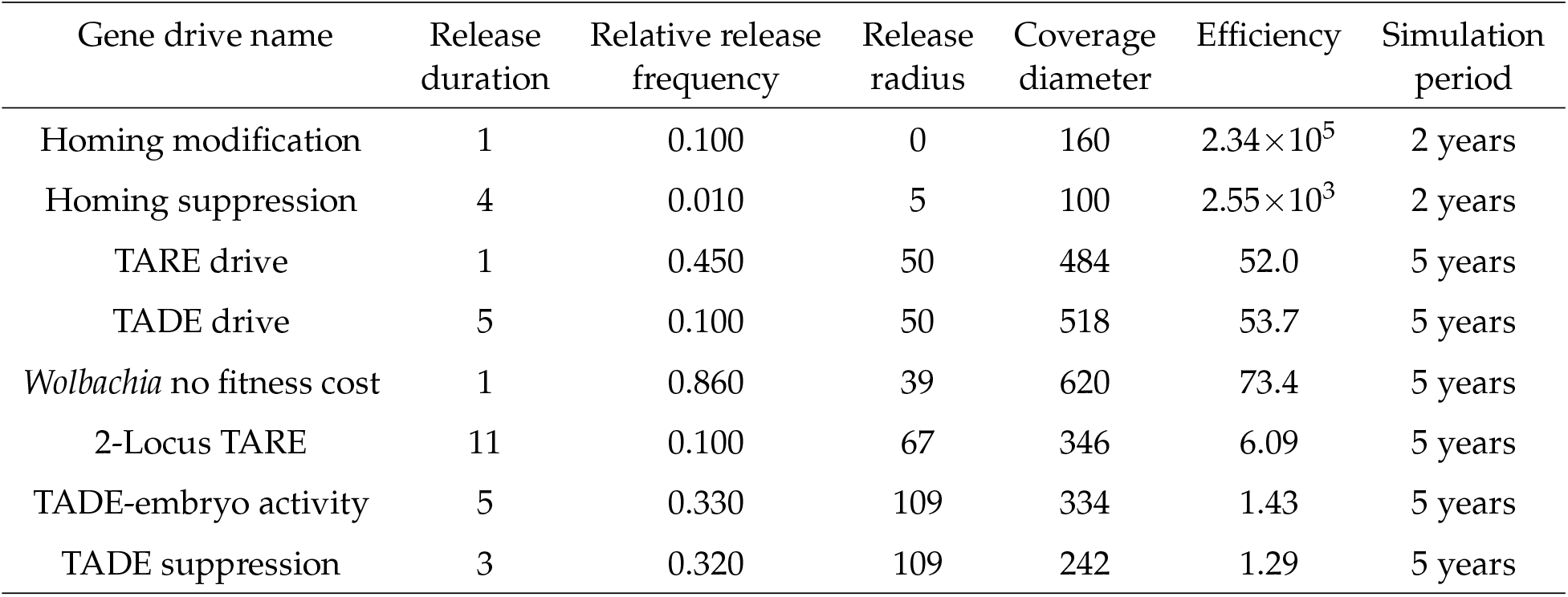
Optimized circular release patterns for different gene drive types.

Despite the far shorter timeframe, homing drives are far more efficient and had far smaller release sizes compared to other systems in this table, showcasing the power of zero-threshold frequency-independent drive. The homing modification drive has a substantial advantage over the suppression drive, which requires a larger and wider release to allow it to form a good drive wave in the timeframe over a wide area, but both systems are extremely efficient. For the frequency-dependent systems, TARE and TADE have good performance in ideal form, spreading widely from their original release area and requiring relatively low release sizes. For greater confinement, 2-locus TARE and *Wolbachia* (due to its fitness cost, otherwise it would be similar to TARE and TADE) still offer a decent level of spread. TADE with embryo activity and TADE suppression are very frequency-dependent and do not offer as high efficiency as the other systems, requiring a wide release. Many of these frequency-dependent systems in particular perform better when releases take place over an extended period of time, preventing loss of effective reproduction by drive individuals at very high concentrations.

### C. Optimal linear drive release

While grid-based circular releases are theoretically the most efficient for area coverage (assuming even release density in the released area), practical applications may sometimes favor releasing along isolated roads if there is not cost-effective access to regions far from roads. We thus also considered scenarios where the drive release took place along a thin line. For our road (or linear) release model, the simulation involves releasing organisms along a path that is one hexagon wide. The key variables adjusted for optimization are just the **release density** and the **release period**, the latter referring to how many generations over which the release takes place. We measured coverage at the end of the simulation and similarly calculated it based on the width of coverage (where drive carrier frequency or population reduction is at least 90%) and density of released individuals per hex of the release area.

In general, results from the optimal linear release (Table 2) were broadly similar to the optimal circle drive release. Efficiency was reduced in most cases because the drive could only spread in two directions from a release point, rather than all directions. For homing modification and for the TARE and TADE drives, efficiency fell by approximately half. However, for the systems with higher thresholds, efficiency loss was more modest. These systems benefited because after initial establishment, the drive was no longer at a geometrical disadvantage (more wild-type outside the circle than drive inside the circle near the circle perimeter). Thus, they only suffered a moderate efficiency loss and in the case of TADE suppression and TADE modification with high embryo activity, actually gained efficiency. It should be noted, though, that this efficiency improvement relies on linear releases with no “ends”, or at least stopping only at a population boundary, such as the edge of an island. A single large release line with an open end may suffer major efficiency loss at the edges and thus be a worse release pattern for high threshold drives than circular releases. Homing suppression drive also slightly gained in efficiency from a linear release compared to a circular release, possibly by establishing a stronger suppression wave more easily from the larger drive release, rather than diffusing over some area without strongly suppressing the population.

**Table 2.** Optimized linear release patterns for different gene drive types.

| Gene drive name | Release duration | Relative release frequency | Coverage diameter | Efficiency | Simulation period |
| --- | --- | --- | --- | --- | --- |
| Homing modification | 1 | 0.01 | 403 | $4.03 \times 10^4$ | 2 years |
| Homing suppression | 1 | 0.01 | 91 | $9.10 \times 10^3$ | 2 years |
| TARE drive | 11 | 1.41 | 443 | 28.6 | 5 years |
| TADE drive | 5 | 4.48 | 521 | 23.3 | 5 years |
| *Wolbachia* no fitness cost | 6 | 3.85 | 595 | 25.8 | 5 years |
| 2-Locus TARE | 18 | 4.76 | 405 | 4.73 | 5 years |
| TADE-embryo activity | 14 | 13.6 | 303 | 1.59 | 5 years |
| TADE suppression | 23 | 4.77 | 173 | 1.65 | 5 years |

### D. Gene drive grid release patterns on Hainan Island

Using optimized parameters from previous models, we implemented a grid release in the Hainan Island model, where each point in the grid was composed of an optimized circular release, arranged in a hexagonal lattice (Figure S4). Because of our 90% coverage area for optimized releases, the locations equidistant between three hexes would likely still reach a high drive frequency in our selected timeframe (over 90% in some of all of the area because of drive influx from multiple directions). Our strategy was to deploy gene drive carriers proportionally to the local adult population at the time of release (week 25 of a year, representing the start of a summer lull in population).

To assess the results, we used two key metrics: area coverage efficiency and population coverage efficiency. Area coverage efficiency is defined as above, while population coverage efficiency is the fraction of humans living in areas with at least 90% drive carrier frequency (modification drives) or 90% mosquito population reduction per drive individual released. Release size itself is now the relative number of newly eclosed adult drive homozygote mosquitoes (or heterozygotes for suppression drives) released. For the homing drives, we assessed a two-year plan, and for other drives, we used a five-year plan for optimization.

Examining release patterns (Figure 4, Table 3), we can see that the homing drives generally had a denser release pattern, but that individual releases within each spot were far smaller than for other drives, in line with our grid-based optimization. Actual drive spread after two years was somewhat spotty for the modification drive, usually high, but often well below 90%, even close to release locations. This is likely because of complexities in the actual population densities near the release area. If the drive is released in a low-density area, it may spread more slowly to surrounding areas that predicted by our earlier optimal release results. This was a reduced issue with suppression drives because its wider wave means that there will be more spillover of drive alleles beyond the predicted radius of population suppression, allowing adjacent drive release areas to work together more effectively to achieve high drive frequency. Frequency-dependent drives demonstrated this pattern even more sharply. For all of them, most area reached nearly 100%, but many regions had near-zero drive frequency, in some cases even within release areas. This is due to influx of wild-type from surrounding higher density areas, which pushed the drive below its threshold (or in the case of the zero-threshold drives, diluted it to such a low level that it lost the ability to spread). These density-dependent effects varied in magnitude for different systems, being particularly apparent in frequency-dependent modification systems.

**Table 3.**
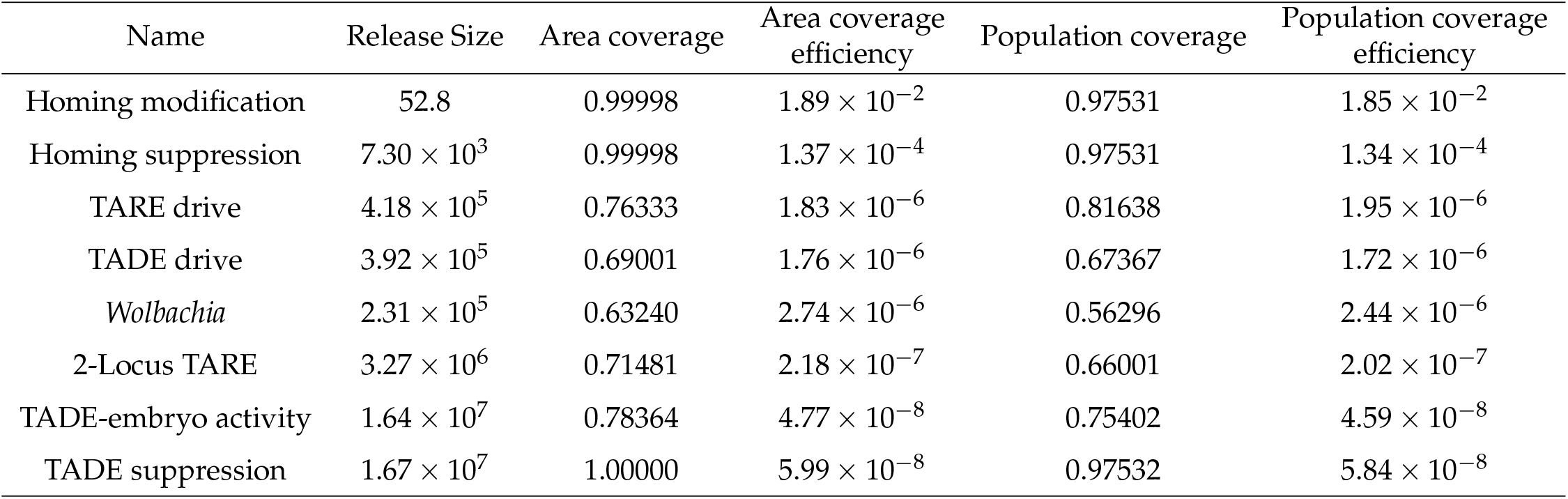
Outcomes of optimized gene drive grid release.

| Name | Release Size | Area coverage | Area coverage efficiency | Population coverage | Population coverage efficiency |
| --- | --- | --- | --- | --- | --- |
| Homing modification | 52.8 | 0.99998 | $1.89 \times 10^{-2}$ | 0.97531 | $1.85 \times 10^{-2}$ |
| Homing suppression | $7.30 \times 10^3$ | 0.99998 | $1.37 \times 10^{-4}$ | 0.97531 | $1.34 \times 10^{-4}$ |
| TARE drive | $4.18 \times 10^5$ | 0.76333 | $1.83 \times 10^{-6}$ | 0.81638 | $1.95 \times 10^{-6}$ |
| TADE drive | $3.92 \times 10^5$ | 0.69001 | $1.76 \times 10^{-6}$ | 0.67367 | $1.72 \times 10^{-6}$ |
| *Wolbachia* | $2.31 \times 10^5$ | 0.63240 | $2.74 \times 10^{-6}$ | 0.56296 | $2.44 \times 10^{-6}$ |
| 2-Locus TARE | $3.27 \times 10^6$ | 0.71481 | $2.18 \times 10^{-7}$ | 0.66001 | $2.02 \times 10^{-7}$ |
| TADE-embryo activity | $1.64 \times 10^7$ | 0.78364 | $4.77 \times 10^{-8}$ | 0.75402 | $4.59 \times 10^{-8}$ |
| TADE suppression | $1.67 \times 10^7$ | 1.00000 | $5.99 \times 10^{-8}$ | 0.97532 | $5.84 \times 10^{-8}$ |

**Figure 4.**
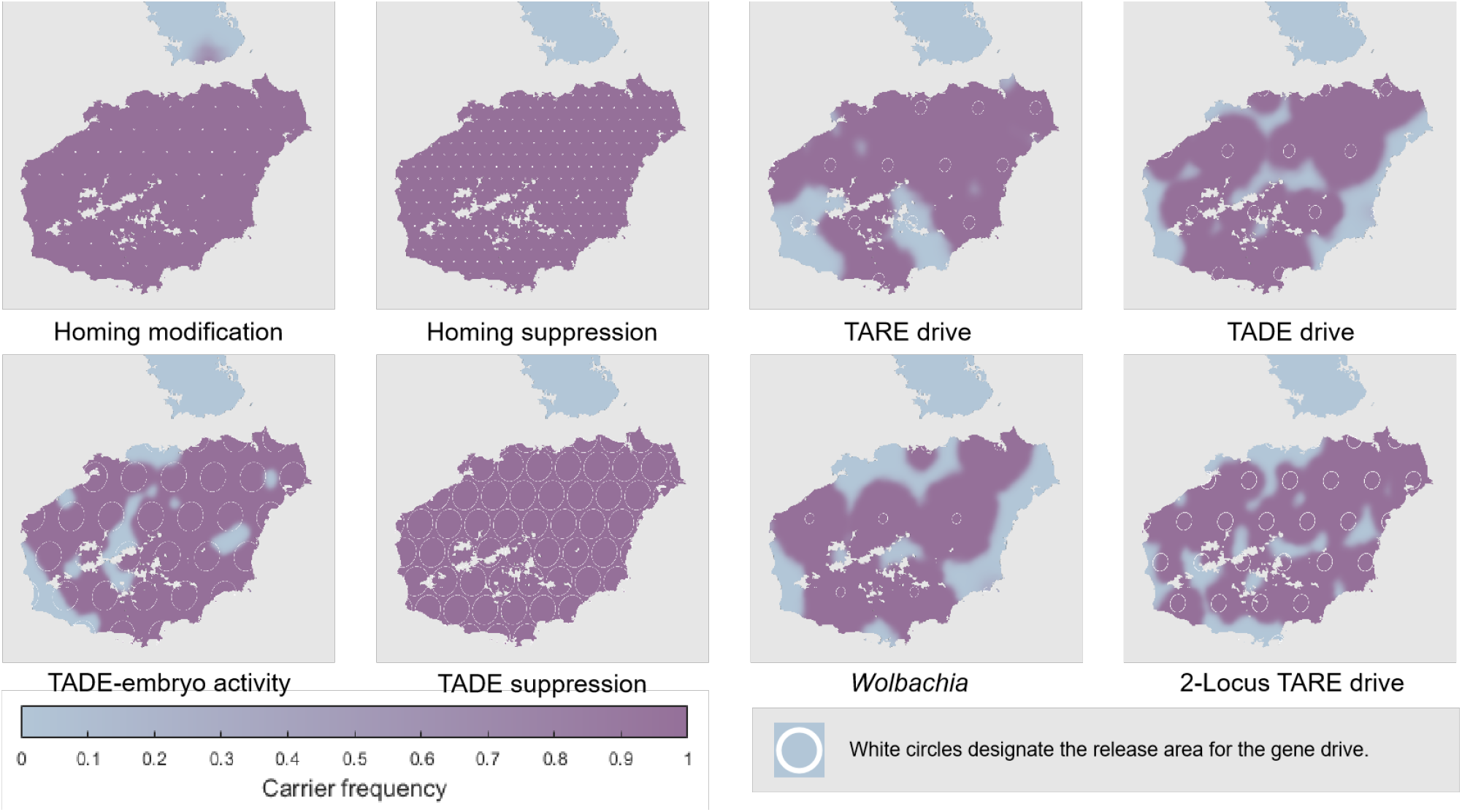
Grid-based release of drive systems on Hainan Island. The map indicates the frequency of the gene drive carriers across Hainan Island following a grid-based release (two years after release for homing drives, five years for others). White circles show where the drive was initially released according to the optimal release patterns in Table 1.

Of all the analyzed drives, only the homing modification drive reached the mainland at significant frequency, though with just a few additional weeks, the homing suppression drive would also spread to high frequency there. Even several of the frequency-dependent but zero-threshold systems did not reach appreciable frequency due to the small absolute level of migration, despite the five-year time span of the simulations.

We also implemented a “linear release” of the drives, utilizing the optimized parameters from our previous work. This was envisioned as a insect release along major roads to minimize costs during the release process. It involved a one-hex-wide release along these roads, except for homing modification, where releases still took place along roads but with intervening hexes with no release (spaced according to optimal distances in the grid release pattern). Efficiency was calculated as above. In general, the releases yielded fairly high coverage of the island (Figure 5), with only homing suppression drive failing to widely suppress the island (though frequencies remained high, and wide suppression would likely have occurred with a slightly greater time interval). Other drives only had a few gaps in areas far from the main roads, which usually also had a low human population density.

**Figure 5.**
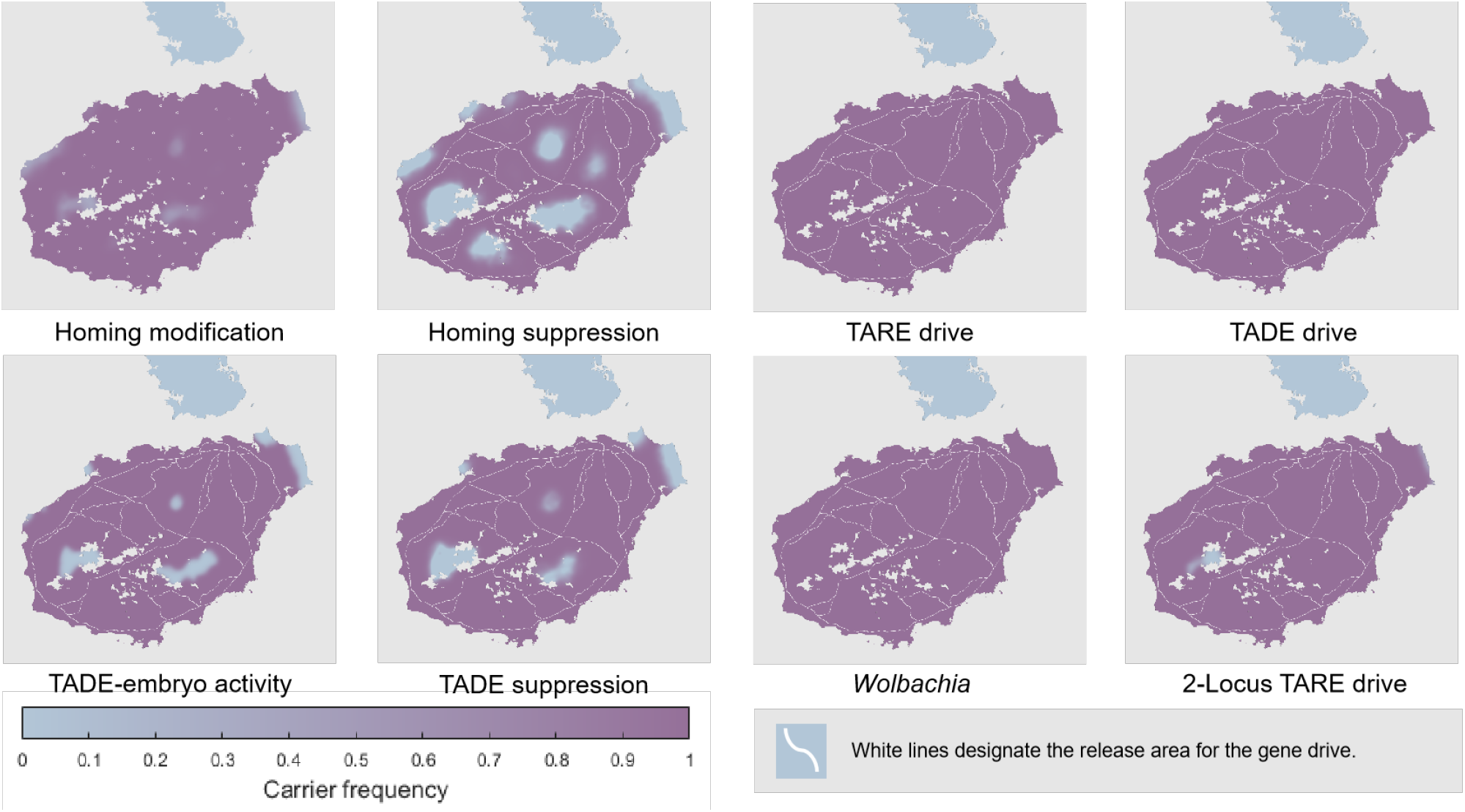
ed release of drive systems on Hainan Island. The map indicates the frequency of the gene drive carriers across Hainan Island following a release along major roads (two years after release for homing drives, five years for others). White lines show where the drive was initially released according to the optimal release patterns in Table 2. Note that the homing modification drive was not released continuously along the whole length of the roads, but at several evenly distributed points along the roads.

**Figure 6.**
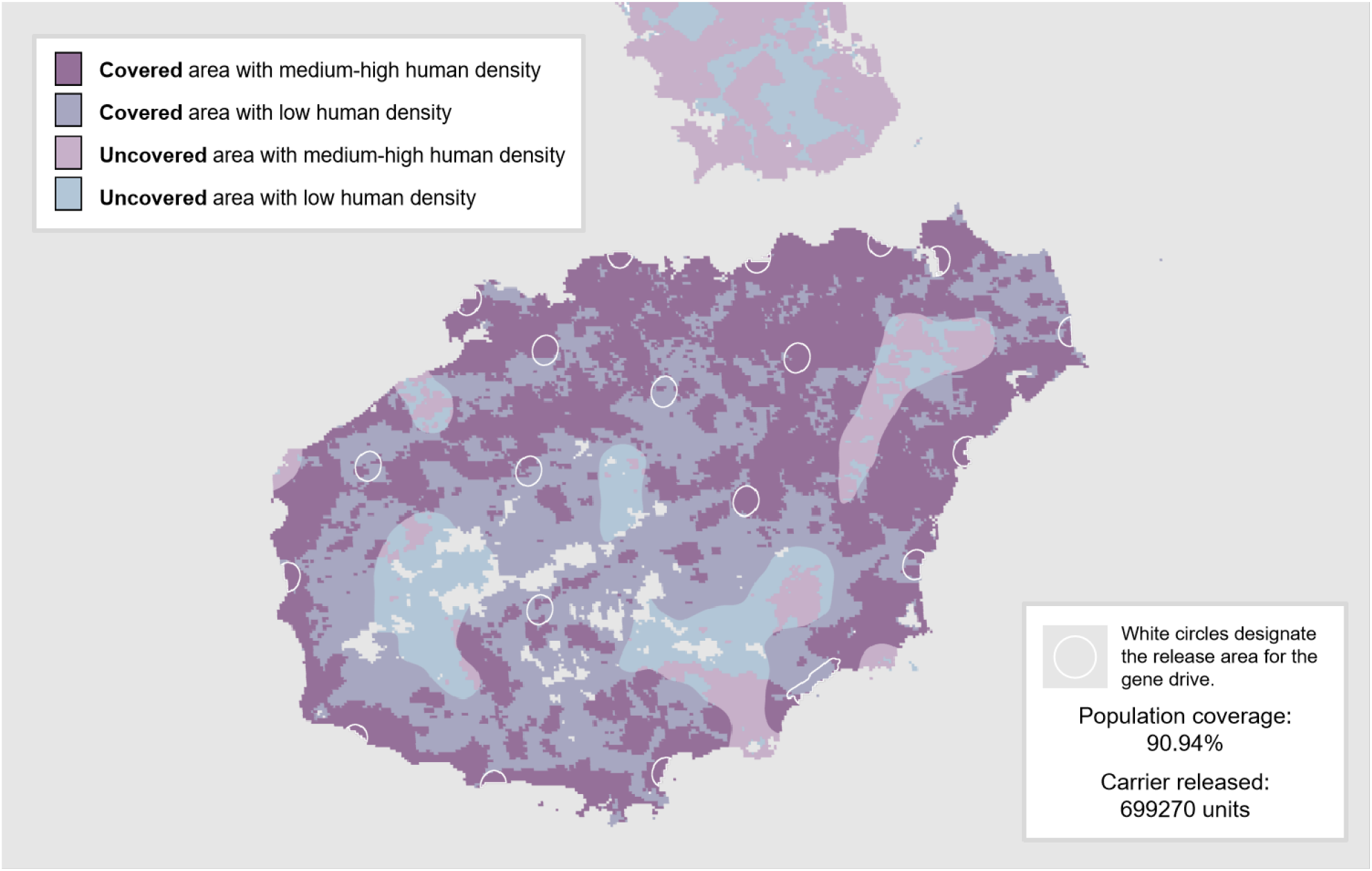
Partially optimized TARE drive release pattern and results. The figure shows the TARE drive release areas with white circles, while also showing final drive coverage with over 90% drive carrier frequency. Populated areas are displayed, and area defined as territory with over 100 people/km^2^ is shown.

Efficiency was generally reduced compared to the optimal grid release (Table 4). For some drives, it was a modest reduction (2-3x), while for some, inefficiency was considerably larger, including homing modification, TADE, and 2-locus TARE. In some cases, this reduced efficiency was caused by two different major roads being nearby, and could have been avoided with a more optimized release. Notably, despite lower efficiency than the grid-release in theory, these ed releases appeared less affected by variations in target population density.

**Table 4.** Outcomes of optimized gene drive release along major roads.

| Name | Release Size | Area coverage | Area coverage efficiency | Population coverage | Population coverage efficiency |
| --- | --- | --- | --- | --- | --- |
| Homing drive | 5.87 | 0.919 | $1.57 \times 10^{-1}$ | 0.943 | $1.61 \times 10^{-1}$ |
| Homing Suppression | $1.70 \times 10^3$ | 0.718 | $4.22 \times 10^{-4}$ | 0.836 | $4.91 \times 10^{-4}$ |
| TARE drive | $2.75 \times 10^6$ | 1.000 | $3.64 \times 10^{-7}$ | 0.975 | $3.55 \times 10^{-7}$ |
| TADE drive | $4.40 \times 10^6$ | 1.000 | $2.27 \times 10^{-7}$ | 0.975 | $2.22 \times 10^{-7}$ |
| *Wolbachia* | $4.52 \times 10^6$ | 1.000 | $2.21 \times 10^{-7}$ | 0.975 | $2.16 \times 10^{-7}$ |
| 2-Locus TARE | $1.81 \times 10^7$ | 0.982 | $5.42 \times 10^{-8}$ | 0.973 | $5.37 \times 10^{-8}$ |
| TADE-embryo activity | $8.20 \times 10^7$ | 0.907 | $1.11 \times 10^{-8}$ | 0.932 | $1.14 \times 10^{-8}$ |
| TADE suppression | $2.25 \times 10^7$ | 0.885 | $3.94 \times 10^{-8}$ | 0.927 | $4.12 \times 10^{-8}$ |

### E. Optimization of a TARE drive release on Hainan Island

While truly optimal releases depend on many factors, such as release capacity, economics of releases, and timeframe, our hex model allows demonstration of a highly efficient release. The regular grid arrangement may lead to inefficiency by causing excessive releases in unpopulated areas and insufficient releases in populated regions, resulting in low coverage and high release numbers. We thus developed a potentially improved release pattern, designed to achieve 90% human coverage with a small release size using TARE, a strong but frequency dependent drive. We started with the evenly spaced grid release and then slightly adjusted release locations to cover areas with high human density. Several releases were also added in coastal areas, where the drive could be protected from geometrical disadvantage (despite smaller release) and where human density was high.

At the end of the fifth year, drive carriers covered 90.9% of Hainan’s population, requiring only 699270 units of drive carrier mosquitoes. This was based on 20 release spots with a total release level of 7 *×* 10^5^

## 4. Discussion

Many different frameworks have been used to assess the outcome of a gene drive release, each with advantages and disadvantages. Here, we developed a new hex-based framework for analyzing gene drive systems. It allows the large scales of patch-based models to be combined with the high fidelity of continuous space models, filling a potentially important niche. By representing the landscape as interconnected hexagonal patches, migration and competition processes are efficiently simulated. This design ensures computational resource savings, enabling users to incorporate a variety of factors such as complex life cycles and heterogeneous terrain.

First, we simulated waves of drive spread, for which we quantified wave speed and shape, demonstrating fairly good agreement with reaction-diffusion models. We thus concluded that with sufficiently high dispersal and an appropriate dispersal kernel, our hex model was capable of mimicking continuous space with high fidelity. To illustrate the capabilities of our hexagonal framework, we then presented two case studies.

First, we used it to determine optimal release patterns for several different gene drives. These release patterns were all constant-density and circular, but could incorporate multiple releases, unlike a previous study that focuses on optimal release shape in space[70]. These optimal release patterns are valuable because they can recommend an actual release strategy that achieves the maximum effect with fixed resources. Other previous studies have focused on critical bubbles, which are closely related to fundamental properties of the system[71], [72], [73], but these not recommended for actual release because they will often not result in drive spread. These optimal releases show that, as expected, some threshold-dependent drives are substantially more efficient than others, though this comes at the cost of lower introduction threshold and consequently less confinement. Homing drives, on the other hand, are vastly more efficient than even frequency-dependent drives that lack a threshold in ideal form. While homing drives are perhaps likely to be further from their “ideal” form than other drives[10], they would still be far more efficient in terms of their spread (the >1000 fold efficiency improvement for homing drives in this study was with a 2-year release plan versus five years for the other drives). Other factors can partly make up for this, but it means that if achieving even moderate drive release numbers may be challenging, use of confined drives should be clearly justified (certainly possible in many situations) if a similar homing drive is available.

Next, we modeled *Culex* mosquitoes with realistic life cycles on Hainan Island, showcasing our framework’s capacity to handle dynamic population density based on seasonally adjusted weather and terrain factors. In this model, we explored optimal gene drive releases on a larger scale, combining several individual locally optimal releases into a whole program to cover the island. Specifically, we analyzed gene drive release in grid and linear patterns. We found that different systems can have substantial variation in how the drive waves combined to fill the intervening territory. In particular, frequency-dependent drive releases should carefully take into account the density of both the release area and surrounding areas. In the future, systematically investigating these for large-scale program optimization may further reduce the necessary resource investment in a genetic control program. We also manually set a release pattern for TARE drive to cover most of the human population for minimal investment, though no doubt this too could be further optimized. Such optimization may be complex, though. Actual releases would have a somewhat fixed production level (though the size of rearing facilities could be determined in advance as a variable), but they could also produce insects for a variable amount of time before the program ends. The total timeframe of the intervention (time to desired results) could also impact the exact release plan. Some parts of this involve economic considerations of both the intervention and cost of the main issue to be addressed, which can be difficult to assess without industrial support. However, our hex-based framework could potentially serve as a basis to explore many scenarios quickly and find programs that are still suitable even if certain parameters are unknown.

With our “linear/road” releases, we also touched on another economic aspect, the cost of insect deployment. This could not only be affected by the location of one or more insect production facilities, but also by the level of connection between the facilities and desired release areas. It is possible that optimal release areas may be poorly served by roads, or even lack roads entirely. This could be a particularly significant issue for the more confined drives in which circular releases tend to have high radii, possibly extending far from secondary roads. This could potentially be overcome by deploying the insects on foot or with off-terrain vehicles, but new drone technology could also be highly suitable in these cases. Yet, even if reasonably effective deployment could be conducted, economic analysis is necessary to determine if it is indeed better than simply using a linear/road release with a greater total number of released insects.

While not a primary component of our model, we briefly considered drive confinement by including actual ferry routes between Hainan and the mainland. Even in ideal form, none of the frequency-dependent drives reached a significant level on the mainland in the five-year timeframe of our model. With zero fitness cost, the zero-threshold ones would have eventually spread to the mainland, but even a small fitness cost would have provided an introduction threshold and prevented this. The unconfined homing drives successfully spread to the mainland, but only the modification drive had sufficient time to reach high frequency there compared to their release location in our short simulations.

While our hex-based model was designed to improve realism, it still has several drawbacks compared to other model systems. First and foremost, the current implementations lacks stochastic variation, which is particularly important in phenomenon such as chasing[22], [25], [74], wherein suppression drives fail to completely eliminate wild-type, despite theoretically having enough drive power. Our mosquito lifecycle model is simplified (such as with weekly timesteps), and our capacity map is also based on a simple model, with modest adjustments from one previously used for a different type of mosquito. The hex-based framework is still composed of discrete patches, so there may be some differences in outcomes compared to a truly continuous model. It also fails to consider potential long-range dispersal by winds or human movement, other than the mainland ferry. Though these are less likely to be important for frequency-dependent drives, they could drastically alter the outcome of a homing drive.

In summary, we used a hex-based framework to assess the performance of several drives and compare some of their properties to reaction-diffusion models. We then examined releases of different drives on Hainan Island as part of a potential *Culex quinquefasciatus* control effort. The efficiency and accuracy of our hex-based framework makes it a good choice for many gene drive modeling applications. Our case scenarios show its potential utility as a decision-support tool. Such tools, especially after further planned improvements, may be valuable for developing specific plans for gene drive releases in the future.

## 5. Acknowledgments

We acknowledge the High-Performance Computing Platform of the Center for Life Science at Peking University for data collection. This study was supported by the Center for Life Sciences and the National Natural Science Foundation of China (grants 32270672 and W2432018).

## 6. Supplemental Information

**Figure S1.**
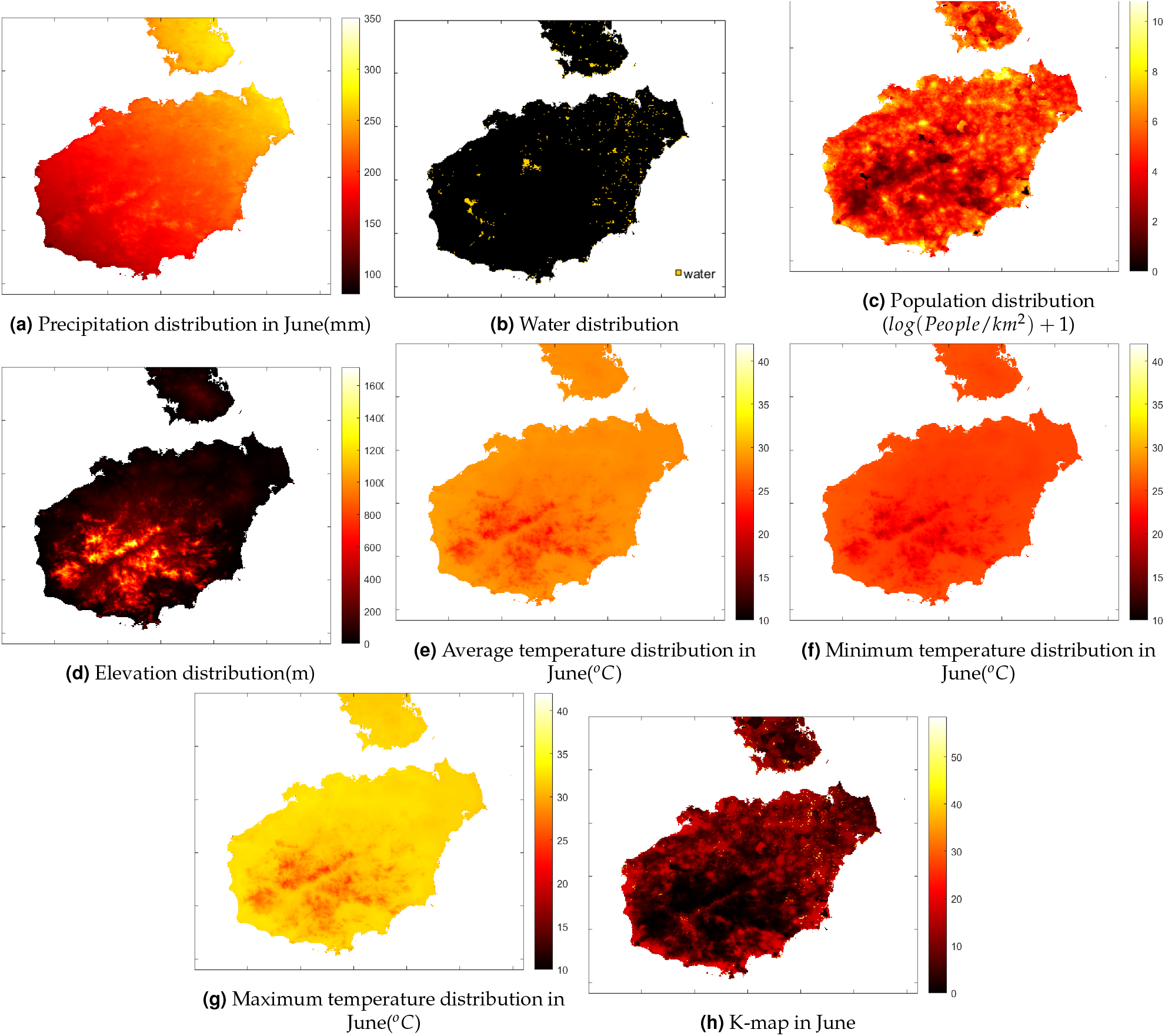
Example characteristics of Hainan Island. Using June as an example, the different figure panels show climate and terrain data used for building the final mosquito carrying capacity map (K-map).

**Figure S2.**
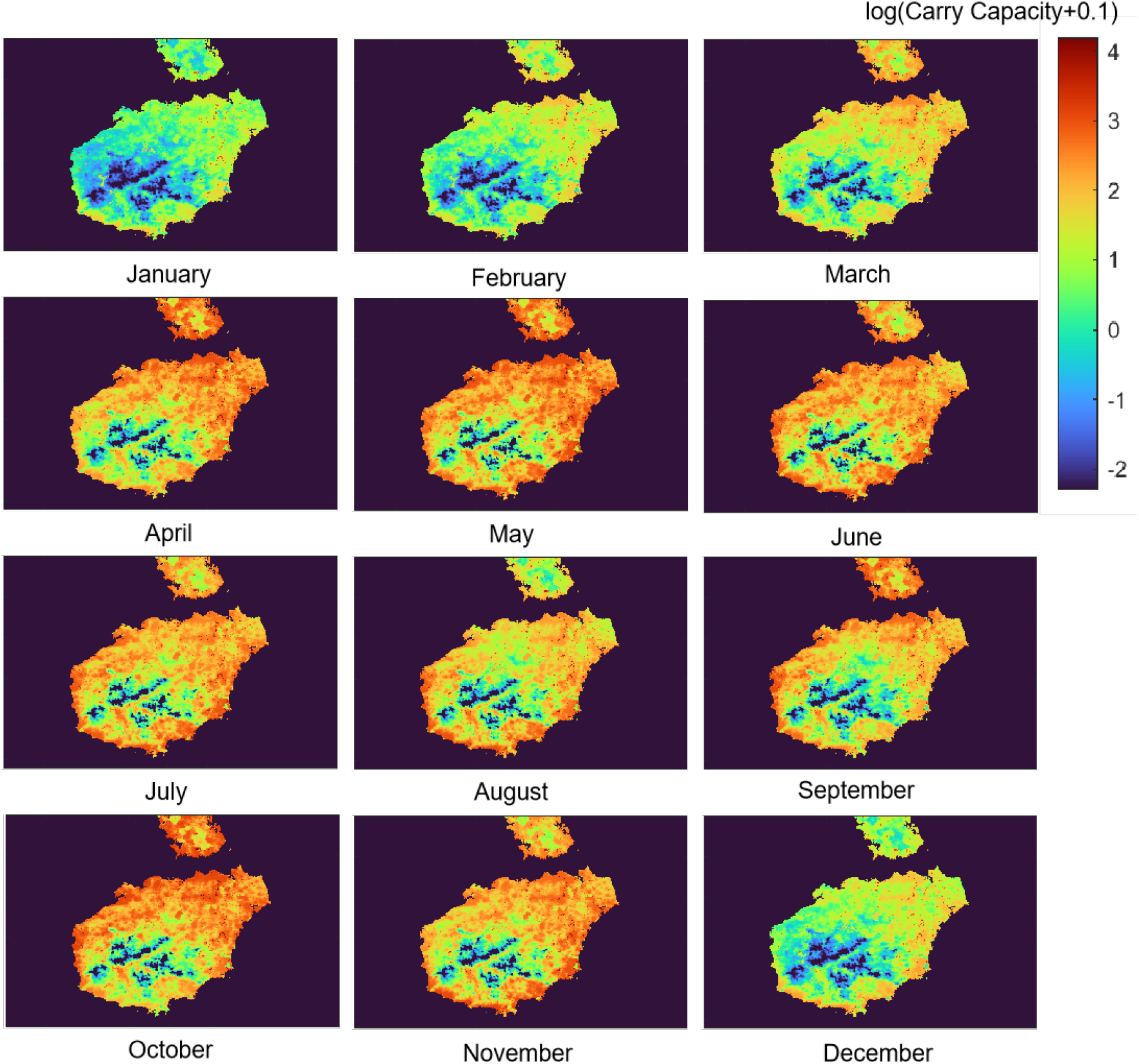
Carrying capacity through time. The figure shows the carrying capacity for each month in the Hainan Island *Culex quinquefasciatus* model.

**Figure S3.**
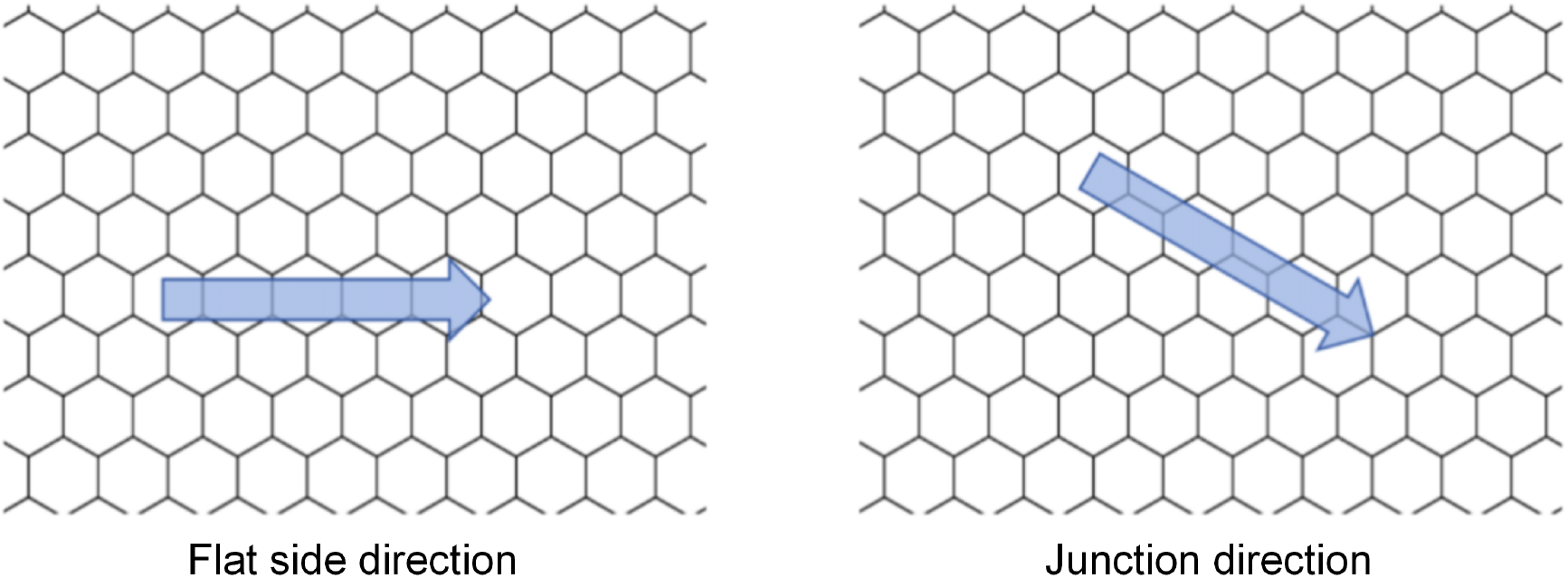
Hex model wave directions. The images show two possible directions for wave advancement through a hex grid.

**Figure S4.**
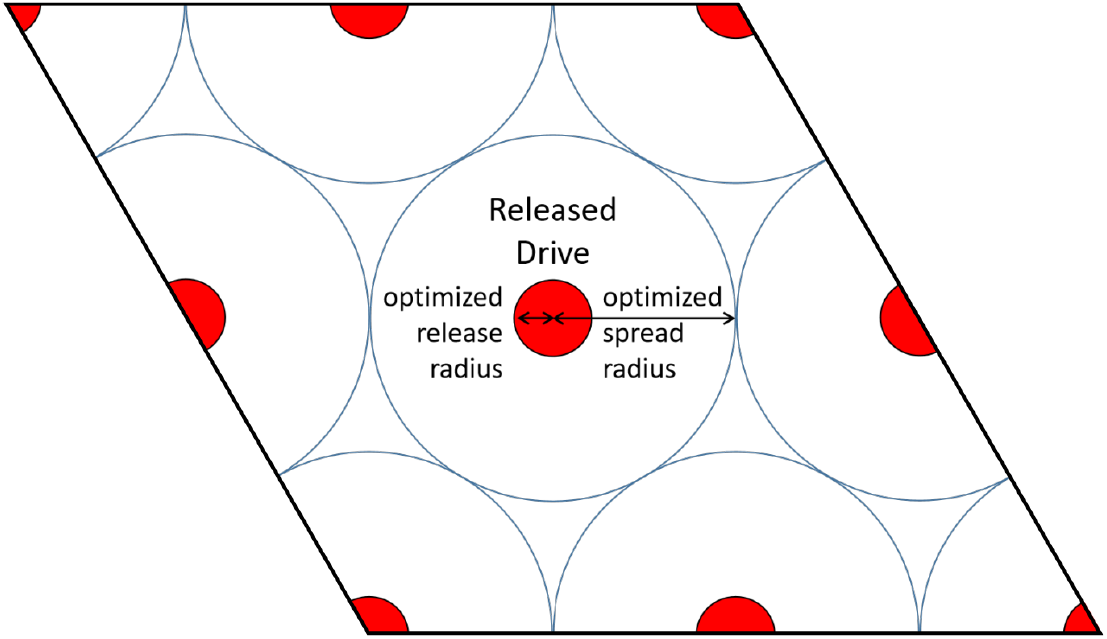
Grid release pattern. The individually optimal release circles are set up in a hexagonal grid over a large areas such that their spread radii touch the six adjacent releases.

**Figure S5.**
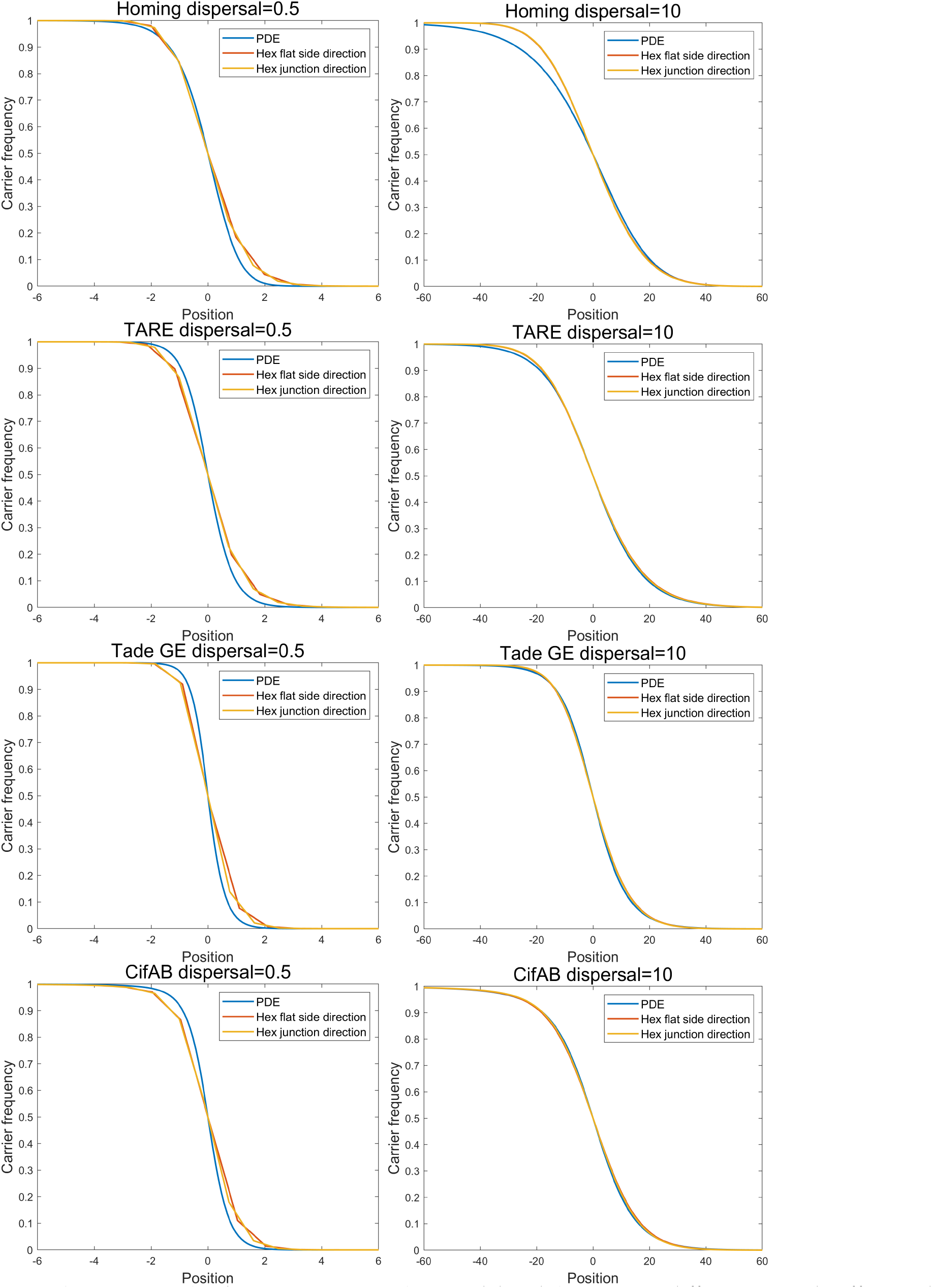
Drive wave shape. Using the discrete-generation hex model and the reaction-diffusion Partial Differential Equation (PDE) model, we show the wave “shape”, specifically the drive carrier frequency as a function of position. We show different wave shape with an average dispersal set to 0.5 or 10 (hexes).

**Figure S6.**
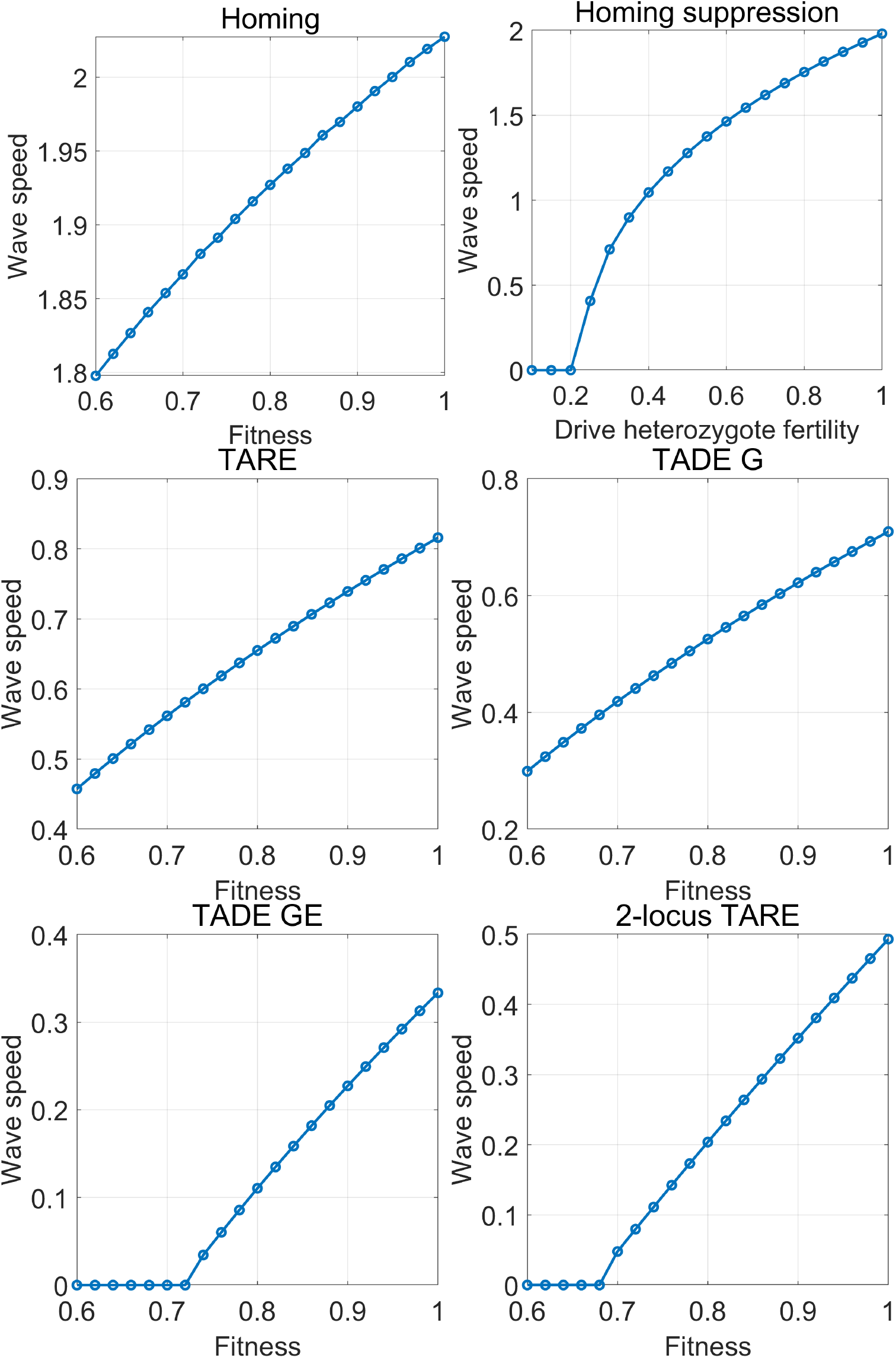
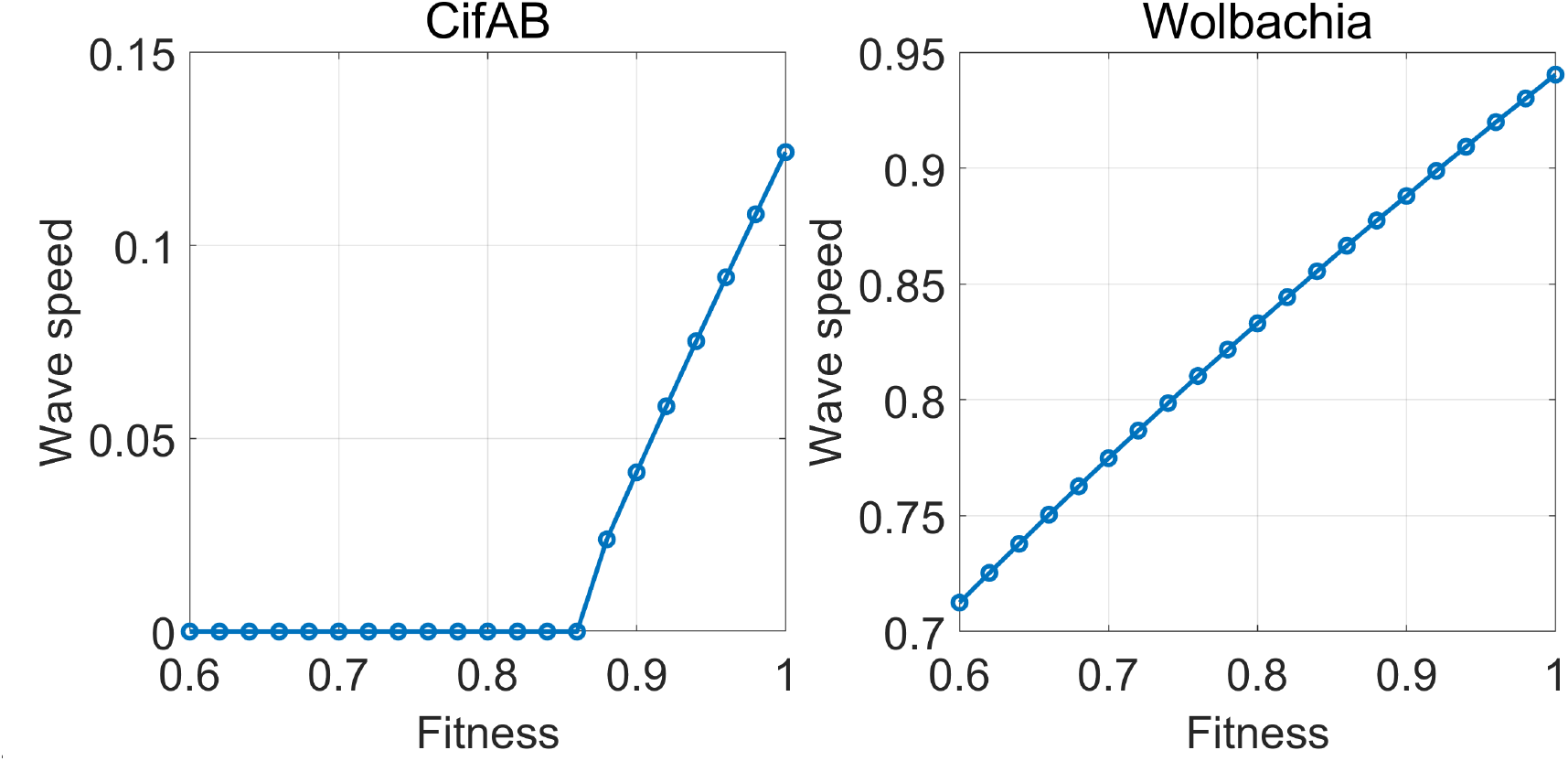
Effect of drive fitness on wave speed. We show the drive wave speed relative to the average dispersal as a function of drive fitness. For modification drives, this is the homozygote fitness, assuming multiplicative per-allele fitness costs. For the homing suppression drive, fitness only affect the fertility of female drive heterozygotes.

**Figure S7.**
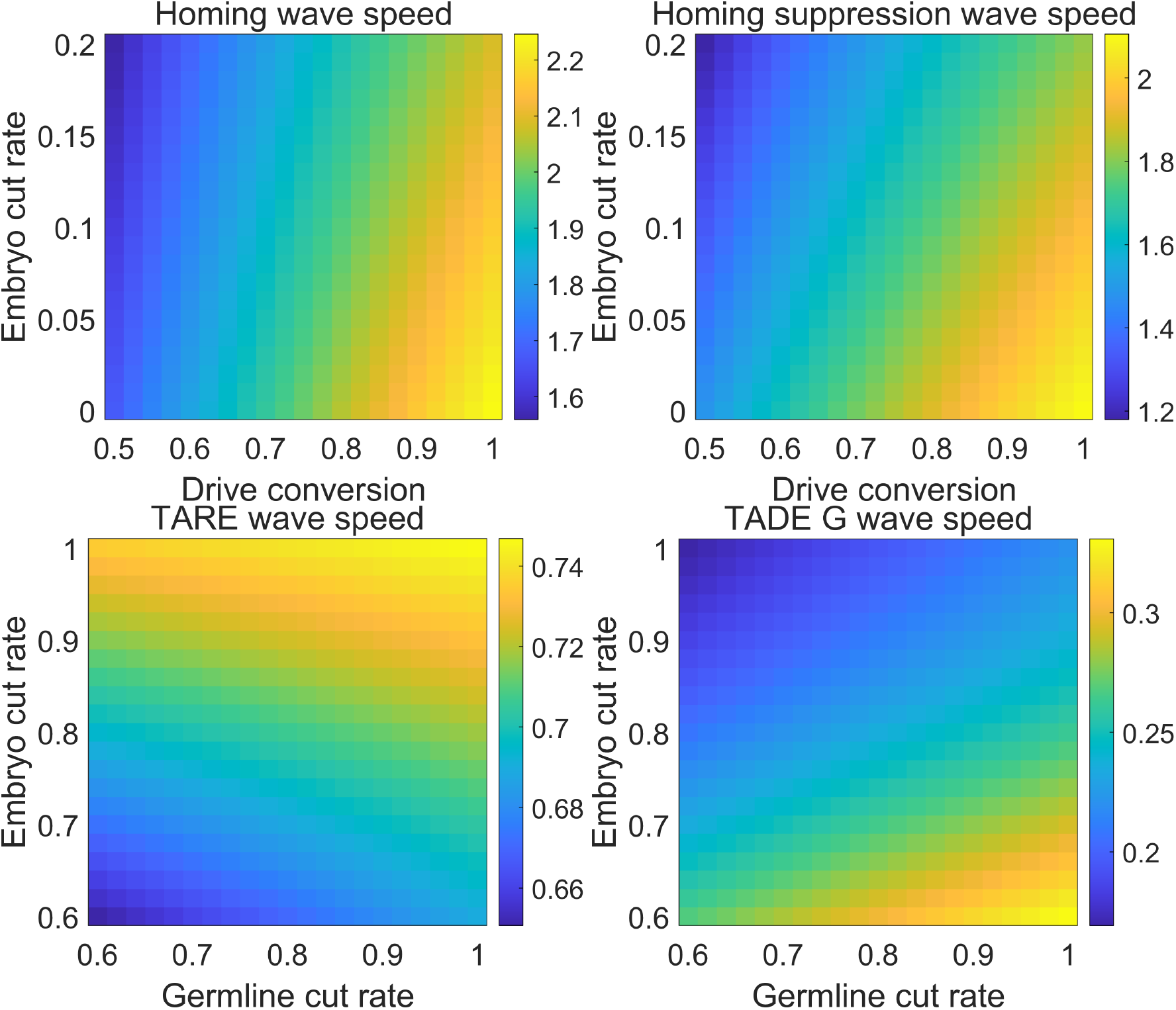
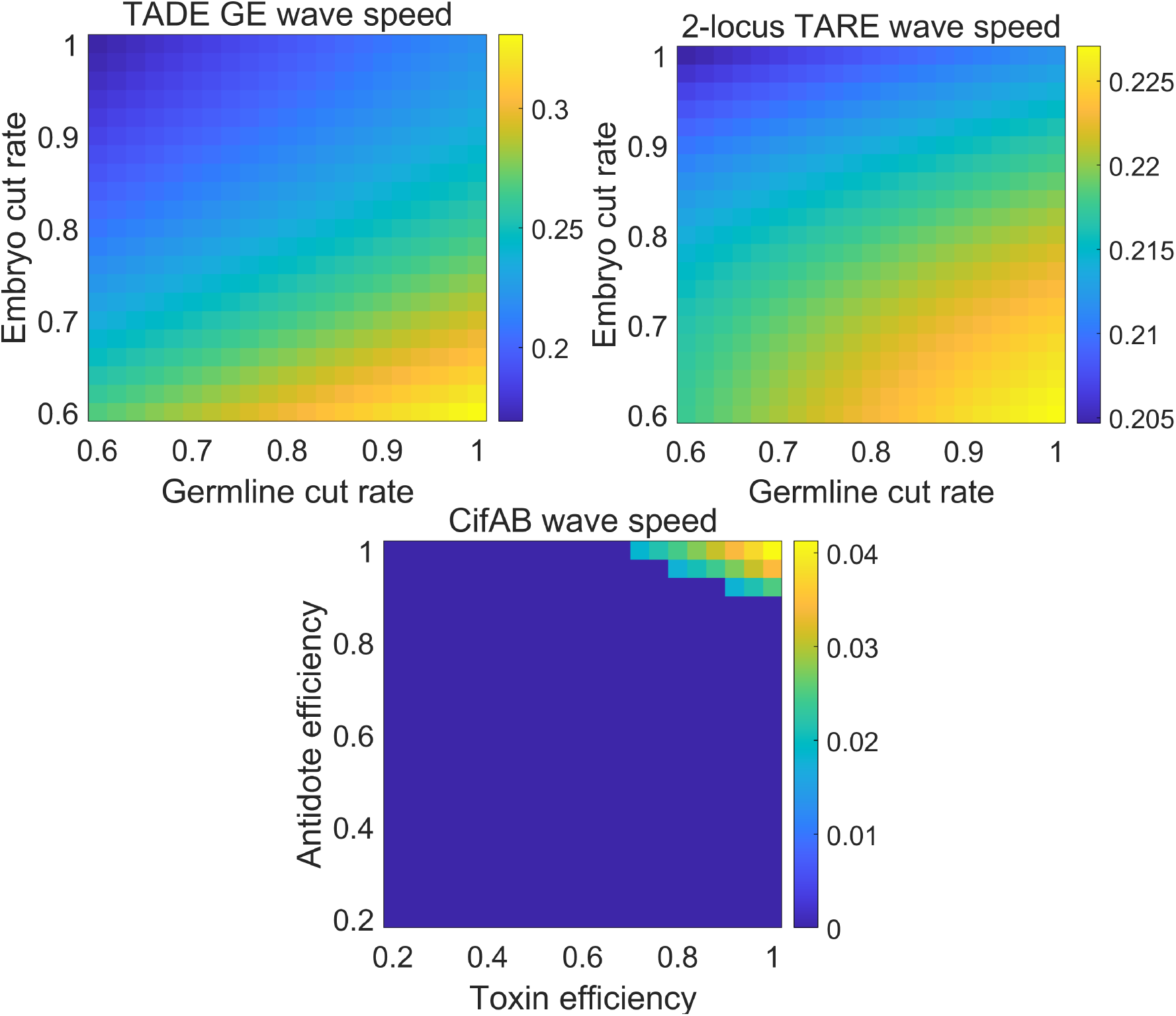
Effect of drive performance on wave speed. We show the drive wave speed relative to the average dispersal as a function of drive performance parameters.

**Figure S8.**
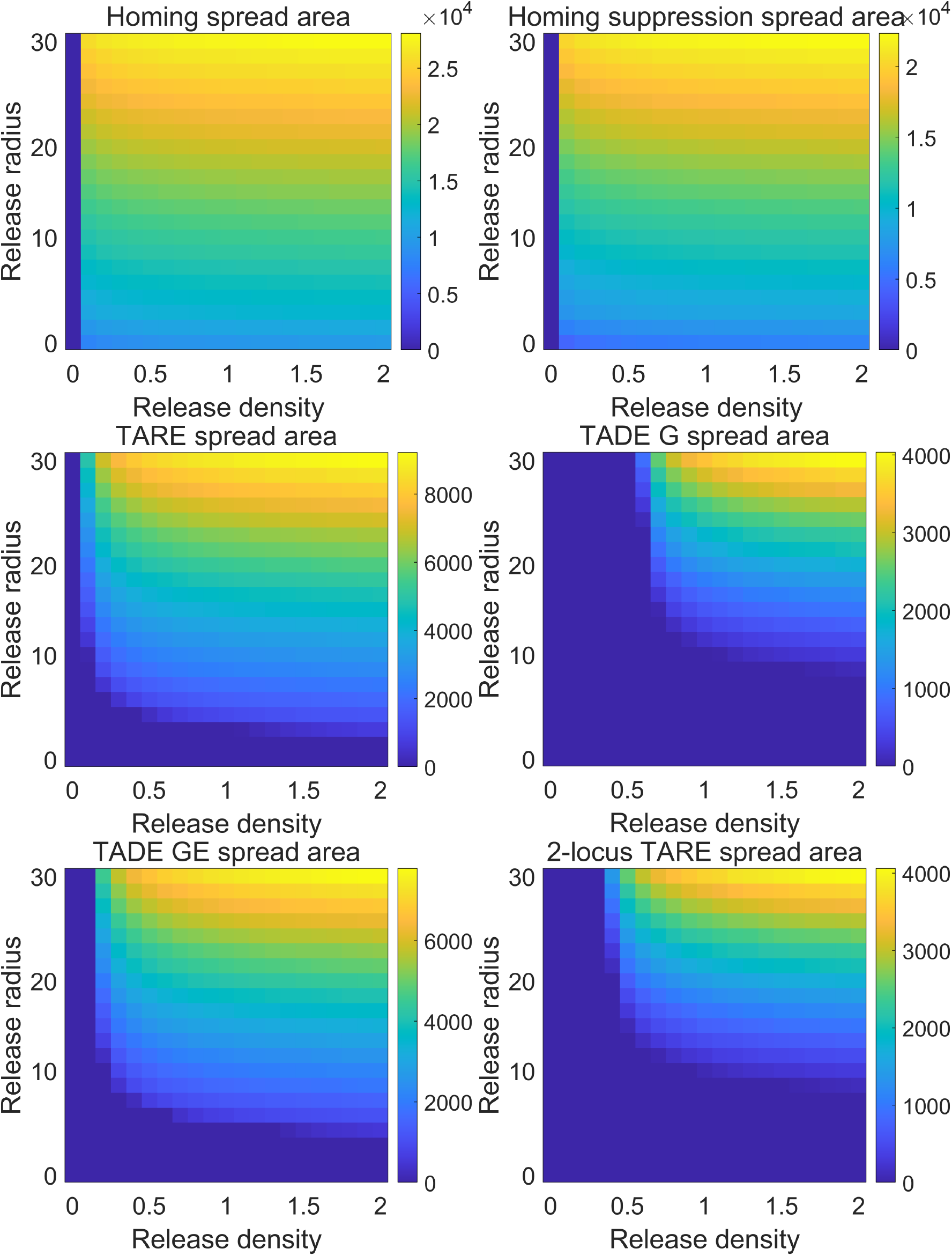
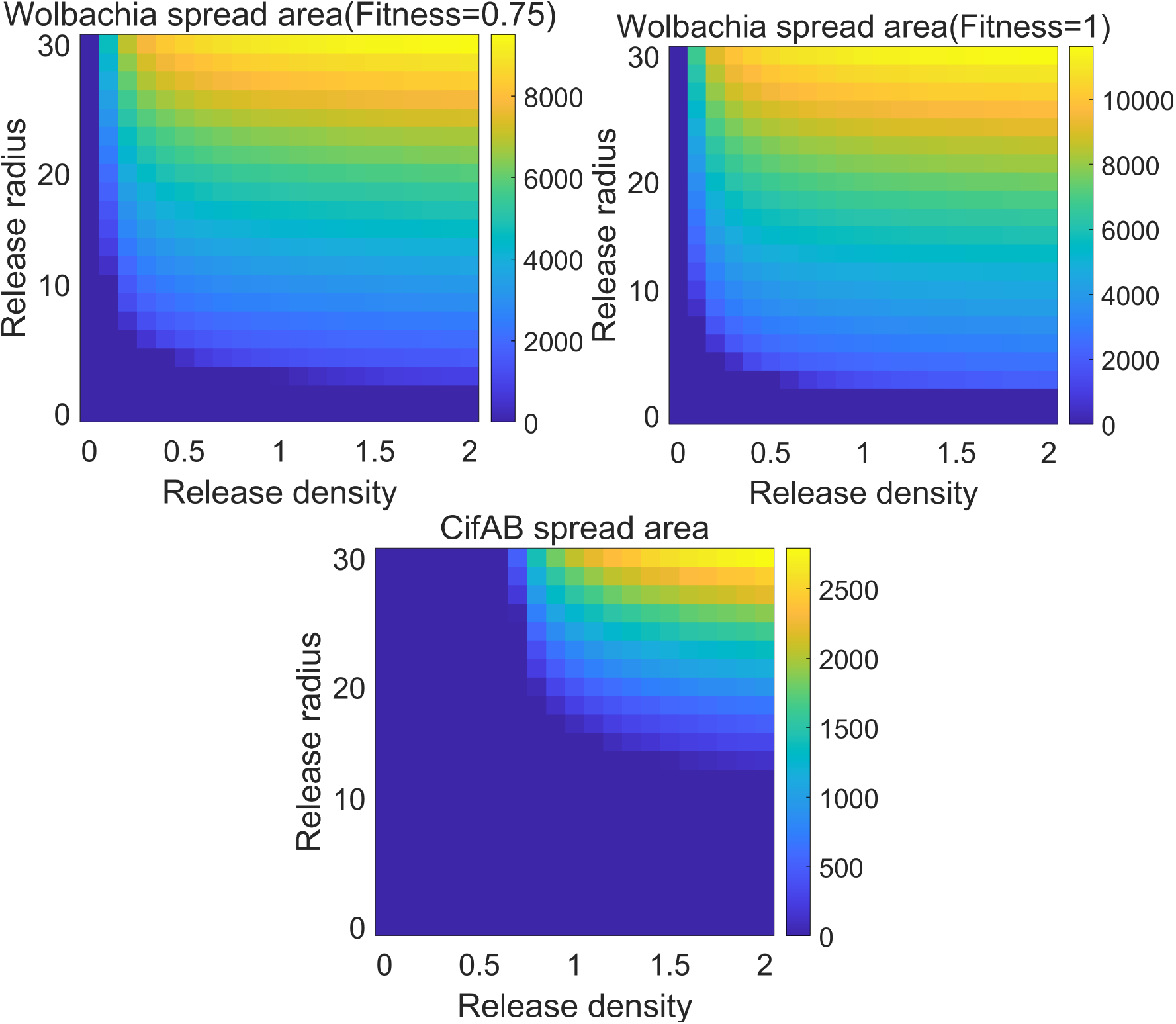
Effect of release pattern on drive spread. In the discrete-generation hex model, we show the number of hexes after two years with at least 50% drive carrier frequency after a release with variable radius and density (relative to the normal population capacity in the release area).

**Figure S9.**
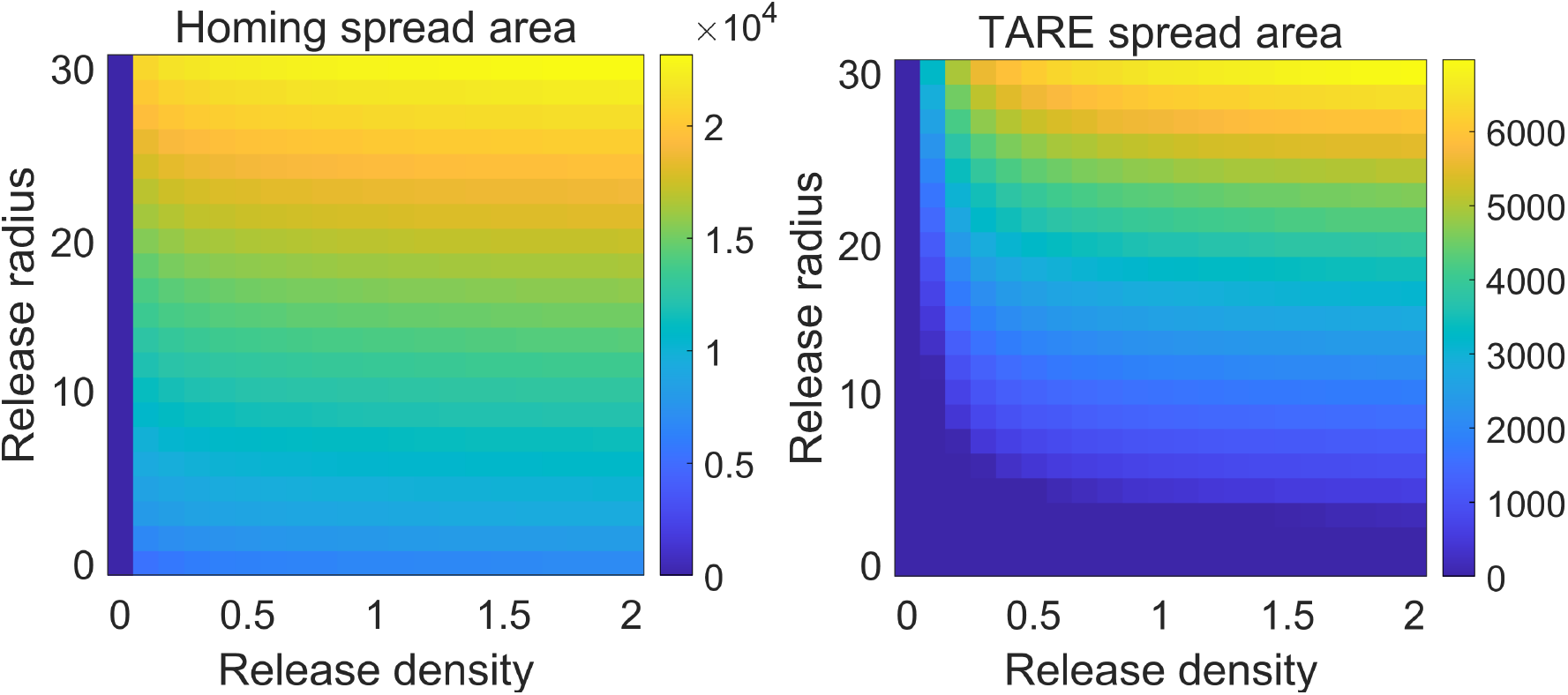
Effect of release pattern on drive spread to 90% drive carrier frequency. As in Figure S8, but showing the number of hexes with at least 90% drive carrier frequency.

## References

[1] B. A. Hay, G. Oberhofer, and M. Guo, “Engineering the composition and fate of wild populations with gene drive,” Annual Review of Entomology, vol. 66, pp. 407–434, 2021.

[2] W. Chen, X. Yang, J. Guo, Y. Han, and J. Champer, “Gene drives and other transgenic approaches for mosquito control,” Trends in Parasitology, 2026.

[3] X. Zhang, Y. Liu, R. Feng, J. Du, and J. Champer, “Engineered gene drives,” in Encyclopedia of Evolutionary Biology (Second Edition), J. B. Wolf and C. A. D. Moraes Russo, Eds., Second Edition, London: Academic Press, 2026, pp. 94–109, ISBN: 978-0-443-15751-6. DOI:10.1016/B978-0-443-15750-9.00138-5 [Online]. Available: https://www.sciencedirect.com/science/article/pii/B9780443157509001385

[4] G.-H. Wang, A. Hoffmann, and J. Champer, “Gene drive and symbiont technologies for control of mosquito-borne diseases,” Annual Review of Entomology, vol. 70, no. Volume 70, 2025, pp. 229–249, 2025.

[5] A. Adolfi, V. M. Gantz, N. Jasinskiene, H.-F. Lee, K. Hwang, G. Terradas, E. A. Bulger, A. Ramaiah, J. B. Bennett, J. Emerson, et al., “Efficient population modification gene-drive rescue system in the malaria mosquito Anopheles stephensi,” Nature communications, vol. 11, no. 1, pp. 1–13, 2020.

[6] J. Champer, E. Yang, E. Lee, J. Liu, A. G. Clark, and P. W. Messer, “A CRISPR homing gene drive targeting a haplolethal gene removes resistance alleles and successfully spreads through a cage population,” Proceedings of the National Academy of Sciences, vol. 117, no. 39, pp. 24377–24383, 2020.

[7] R. Carballar-Lejarazú, Y. Dong, T. B. Pham, T. Tushar, R. M. Corder, A. Mondal, H. M. Sánchez C, H.-F. Lee, J. M. Marshall, G. Dimopoulos, et al., “Dual effector population modification gene-drive strains of the african malaria mosquitoes, anopheles gambiae and anopheles coluzzii,” Proceedings of the National Academy of Sciences, vol. 120, no. 29, e2221118120, 2023.

[8] K. Kyrou, A. M. Hammond, R. Galizi, N. Kranjc, A. Burt, A. K. Beaghton, T. Nolan, and A. Crisanti, “A CRISPR–Cas9 gene drive targeting doublesex causes complete population suppression in caged Anopheles gambiae mosquitoes,” Nature Biotechnology, vol. 36, no. 11, pp. 1062–1066, 2018.

[9] J. Du, W. Chen, X. Jia, X. Xu, E. Yang, R. Zhou, Y. Zhang, M. Metzloff, P. W. Messer, and J. Champer, “Germline Cas9 promoters with improved performance for homing gene drive,” Nature Communications, vol. 15, no. 1, p. 4560, 2024.

[10] J. Champer, I. K. Kim, S. E. Champer, A. G. Clark, and P. W. Messer, “Performance analysis of novel toxin-antidote CRISPR gene drive systems,” BMC biology, vol. 18, no. 1, pp. 1–17, 2020.

[11] J. Champer, S. E. Champer, I. K. Kim, A. G. Clark, and P. W. Messer, “Design and analysis of CRISPR-based underdominance toxin-antidote gene drives,” Evolutionary Applications, vol. 14, no. 4, pp. 1052–1069, 2021.

[12] S. Dhole, M. R. Vella, A. L. Lloyd, and F. Gould, “Invasion and migration of spatially self-limiting gene drives: a comparative analysis,” Evolutionary applications, vol. 11, no. 5, pp. 794–808, 2018.

[13] M. P. Edgington and L. S. Alphey, “Conditions for success of engineered underdominance gene drive systems,” Journal of theoretical biology, vol. 430, pp. 128–140, 2017.

[14] D. Khamis, C. El Mouden, K. Kura, and M. B. Bonsall, “Ecological effects on underdominance threshold drives for vector control,” Journal of Theoretical Biology, vol. 456, pp. 1–15, 2018.

[15] J. Li and J. Champer, “Harnessing Wolbachia cytoplasmic incompatibility alleles for confined gene drive: A modeling study,” PLoS Genetics, vol. 19, no. 1, e1010591, 2023.

[16] J. Champer, E. Lee, E. Yang, C. Liu, A. G. Clark, and P. W. Messer, “A toxin-antidote CRISPR gene drive system for regional population modification,” Nature Communications, vol. 11, no. 1, pp. 1–10, 2020.

[17] M. Maselko, N. Feltman, A. Upadhyay, A. Hayward, S. Das, N. Myslicki, A. J. Peterson, M. B. O’Connor, and M. J. Smanski, “Engineering multiple species-like genetic incompatibilities in insects,” Nature communications, vol. 11, no. 1, pp. 1–7, 2020.

[18] G. Oberhofer, T. Ivy, and B. A. Hay, “Cleave and rescue, a Novel Selfish Genetic Element and General Strategy for Gene Drive,” Proceedings of the National Academy of Sciences, vol. 116, no. 13, pp. 6250–6259, 2019.

[19] R. G. Reeves, J. Bryk, P. M. Altrock, J. A. Denton, and F. A. Reed, “First steps towards underdominant genetic transformation of insect populations,” PLoS One, vol. 9, no. 5, e97557, 2014.

[20] K. W. Okamoto, M. A. Robert, A. L. Lloyd, and F. Gould, “A reduce and replace strategy for suppressing vector-borne diseases: Insights from a stochastic, spatial model,” PLOS ONE, vol. 8, pp. 1–16, Dec. 2013. DOI:10.1371/journal.pone.0081860 [Online]. Available: https://doi.org/10.1371/journal.pone.0081860

[21] M. Pan and J. Champer, “Making waves: Comparative analysis of gene drive spread characteristics in a continuous space model,” Molecular Ecology, vol. 32, no. 20, pp. 5673–5694, 2023.

[22] Y. Liu and J. Champer, “Modelling homing suppression gene drive in haplodiploid organisms,” Proceedings of the Royal Society B, vol. 289, no. 1972, p. 20220320, 2022.

[23] Y. Liu, W. Teo, H. Yang, and J. Champer, “Adversarial interspecies relationships facilitate population suppression by gene drive in spatially explicit models,” Ecology Letters, vol. 26, no. 7, pp. 1174–1185, 2023.

[24] S. E. Champer, N. Oakes, R. Sharma, P. García-Díaz, J. Champer, and P. W. Messer, “Modeling crispr gene drives for suppression of invasive rodents using a supervised machine learning framework,” PLoS computational biology, vol. 17, no. 12, e1009660, 2021.

[25] J. Champer, I. K. Kim, S. E. Champer, A. G. Clark, and P. W. Messer, “Suppression gene drive in continuous space can result in unstable persistence of both drive and wild-type alleles,” Molecular Ecology, vol. 30, no. 4, pp. 1086–1101, 2021.

[26] Y. Liu, S. E. Champer, B. C. Haller, and J. Champer, “Modeling control of invasive fire ants by gene drive,” Advanced Science, vol. 12, no. 46, e04653, 2025.

[27] S. E. Champer, B. Chae, B. C. Haller, J. Champer, and P. W. Messer, “Resource-explicit interactions in spatial population models,” Methods in ecology and evolution, vol. 15, no. 12, pp. 2316–2330, 2024.

[28] J. S. Marshall, “Discrete-element modeling of particulate aerosol flows,” Journal of Computational Physics, vol. 228, no. 5, pp. 1541–1561, 2009.

[29] A. R. North, A. Burt, and H. C. J. Godfray, “Modelling the potential of genetic control of malaria mosquitoes at national scale,” BMC biology, vol. 17, pp. 1–12, 2019.

[30] C. J. Battey, P. L. Ralph, and A. D. Kern, “Space is the place: Effects of continuous spatial structure on analysis of population genetic data,” Genetics, vol. 215, no. 1, pp. 193–214, 2020.

[31] E. Lundgren and P. L. Ralph, “Are populations like a circuit? comparing isolation by resistance to a new coalescent-based method,” Molecular ecology resources, vol. 19, no. 6, pp. 1388–1406, 2019.

[32] C. P. Birch, S. P. Oom, and J. A. Beecham, “Rectangular and hexagonal grids used for observation, experiment and simulation in ecology,” Ecological modelling, vol. 206, no. 3-4, pp. 347–359, 2007.

[33] J. M. Marshall and B. A. Hay, “Confinement of gene drive systems to local populations: a comparative analysis,” Journal of Theoretical Biology, vol. 294, pp. 153–171, 2012.

[34] H. M. Sánchez C, J. B. Bennett, S. L. Wu, G. Rašić, O. S. Akbari, and J. M. Marshall, “Modeling confinement and reversibility of threshold-dependent gene drive systems in spatially-explicit Aedes aegypti populations,” BMC biology, vol. 18, pp. 1–14, 2020.

[35] R. Geci, K. Willis, and A. Burt, “Gene drive designs for efficient and localisable population suppression using Y-linked editors,” PLoS Genetics, vol. 18, no. 12, e1010550, 2022.

[36] P. A. Hancock, A. North, A. W. Leach, P. Winskill, A. C. Ghani, H. C. J. Godfray, A. Burt, and J. D. Mumford, “The potential of gene drives in malaria vector species to control malaria in african environments,” Nature Communications, vol. 15, no. 1, p. 8976, 2024.

[37] A. Beaghton, P. J. Beaghton, and A. Burt, “Gene drive through a landscape: Reaction–diffusion models of population suppression and elimination by a sex ratio distorter,” Theoretical population biology, vol. 108, pp. 51–69, 2016.

[38] S. Zhang and J. Champer, “Performance characteristics allow for confinement of a CRISPR toxin– antidote gene drive for population suppression in a reaction–diffusion model,” Proceedings B, vol. 291, no. 2025, p. 20240500, 2024.

[39] L. Girardin and F. Débarre, “Demographic feedbacks can hamper the spatial spread of a gene drive,” Journal of Mathematical Biology, vol. 83, no. 6, p. 67, 2021.

[40] L. Girardin, V. Calvez, and F. Débarre, “Catch me if you can: A spatial model for a brake-driven gene drive reversal,” Bulletin of Mathematical Biology, vol. 81, no. 12, pp. 5054–5088, 2019.

[41] H. Tanaka, H. A. Stone, and D. R. Nelson, “Spatial gene drives and pushed genetic waves,” Proceedings of the National Academy of Sciences, vol. 114, no. 32, pp. 8452–8457, 2017.

[42] N. H. Schumaker and A. Brookes, “Hexsim: A modeling environment for ecology and conservation,” Landscape Ecology, vol. 33, pp. 197–211, 2018.

[43] C. Liao, T. Zhou, D. Xu, R. Barnes, G. Bisht, H.-Y. Li, Z. Tan, T. Tesfa, Z. Duan, D. Engwirda, et al., “Advances in hexagon mesh-based flow direction modeling,” Advances in Water Resources, vol. 160, p. 104099, 2022.

[44] J. L. Holland and G. D. Gottfredson, “Studies of the hexagonal model: An evaluation (or, the perils of stalking the perfect hexagon),” Journal of vocational behavior, vol. 40, no. 2, pp. 158–170, 1992.

[45] D. P. Petersen and D. Middleton, “Sampling and reconstruction of wave-number-limited functions in n-dimensional euclidean spaces,” Information and control, vol. 5, no. 4, pp. 279–323, 1962.

[46] R. M. Mersereau, “The processing of hexagonally sampled two-dimensional signals,” Proceedings of the IEEE, vol. 67, no. 6, pp. 930–949, 1979.

[47] D. E. Staunton, V. J. Merluzzi, R. Rothlein, R. Barton, S. D. Marlin, and T. A. Springer, “A cell adhesion molecule, icam-1, is the major surface receptor for rhinoviruses,” Cell, vol. 56, no. 5, pp. 849–853, 1989.

[48] G. F. Davies, “Geophysical and isotopic constraints on mantle convection: An interim synthesis,” Journal of Geophysical Research: Solid Earth, vol. 89, no. B7, pp. 6017–6040, 1984.

[49] A. Rosenfeld, “Connectivity in digital pictures,” Journal of the ACM (JACM), vol. 17, no. 1, pp. 146–160, 1970.

[50] S. A. Coleman, B. W. Scotney, and S. Suganthan, “Feature extraction on range images-a new approach,” in Proceedings 2007 IEEE International Conference on Robotics and Automation, IEEE, 2007, pp. 1098–1103.

[51] E. Bier, “Gene drives gaining speed,” Nature Reviews Genetics, vol. 23, no. 1, pp. 5–22, 2022.

[52] S. E. Champer, S. Y. Oh, C. Liu, Z. Wen, A. G. Clark, P. W. Messer, and J. Champer, “Computational and experimental performance of CRISPR homing gene drive strategies with multiplexed gRNAs,” Science Advances, vol. 6, no. 10, eaaz0525, 2020.

[53] S. Hou, J. Chen, R. Feng, X. Xu, N. Liang, and J. Champer, “A homing rescue gene drive with multiplexed gRNAs reaches high frequency in cage populations but generates functional resistance,” Journal of Genetics and Genomics, 2024.

[54] E. Yang, M. Metzloff, A. M. Langmüller, X. Xu, A. G. Clark, P. W. Messer, and J. Champer, “A homing suppression gene drive with multiplexed gRNAs maintains high drive conversion efficiency and avoids functional resistance alleles,” G3, vol. 12, no. 6, jkac081, 2022.

[55] E. I. Green, E. Jaouen, D. Klug, R. P. Olmo, A. Gautier, S. Blandin, and E. Marois, “A population modification gene drive targeting both saglin and lipophorin impairs plasmodium transmission in anopheles mosquitoes,” Elife, vol. 12, e93142, 2023.

[56] M. A. Anderson, E. Gonzalez, M. P. Edgington, J. X. Ang, D.-K. Purusothaman, L. Shack-leford, K. Nevard, S. A. Verkuijl, T. Harvey-Samuel, P. T. Leftwich, et al., “A multiplexed, confinable crispr/cas9 gene drive can propagate in caged aedes aegypti populations,” Nature Communications, vol. 15, no. 1, p. 729, 2024.

[57] T. A. Prowse, P. Cassey, J. V. Ross, C. Pfitzner, T. A. Wittmann, and P. Thomas, “Dodging silver bullets: Good crispr gene-drive design is critical for eradicating exotic vertebrates,” Proceedings of the Royal Society B: Biological Sciences, vol. 284, no. 1860, p. 20170799, 2017.

[58] I. Morianou, L. Phillimore, B. S. Khatri, L. Marston, M. Gribble, A. Burt, F. Bernardini, A. M. Hammond, T. Nolan, and A. Crisanti, “Engineering resilient gene drives towards sustainable malaria control: Predicting, testing and overcoming target site resistance,” bioRxiv, pp. 2024–10, 2024.

[59] Y. Zhu and J. Champer, “Simulations reveal high efficiency and confinement of a population suppression crispr toxin-antidote gene drive,” ACS Synthetic Biology, vol. 12, no. 3, pp. 809–819, 2023.

[60] A. A. Hoffmann, B. Montgomery, J. Popovici, I. Iturbe-Ormaetxe, P. Johnson, F. Muzzi, M. Greenfield, M. Durkan, Y. Leong, Y. Dong, et al., “Successful establishment of Wolbachia in Aedes populations to suppress dengue transmission,” Nature, vol. 476, no. 7361, pp. 454–457, 2011.

[61] R. Kaur, J. D. Shropshire, K. L. Cross, B. Leigh, A. J. Mansueto, V. Stewart, S. R. Bordenstein, and S. R. Bordenstein, “Living in the endosymbiotic world of Wolbachia: a centennial review,” Cell Host & Microbe, vol. 29, no. 6, pp. 879–893, 2021.

[62] P. A. Hancock, S. A. Ritchie, C. J. Koenraadt, T. W. Scott, A. A. Hoffmann, and H. C. J. Godfray, “Predicting the spatial dynamics of wolbachia infections in aedes aegypti arbovirus vector populations in heterogeneous landscapes,” Journal of Applied Ecology, vol. 56, no. 7, pp. 1674–1686, 2019.

[63] S. E. Champer, I. K. Kim, A. G. Clark, P. W. Messer, and J. Champer, “Anopheles homing suppression drive candidates exhibit unexpected performance differences in simulations with spatial structure,” Elife, vol. 11, e79121, 2022.

[64] A. T. Ciota, A. C. Matacchiero, A. M. Kilpatrick, and L. D. Kramer, “The effect of temperature on life history traits of culex mosquitoes,” Journal of medical entomology, vol. 51, no. 1, pp. 55–62, 2014.

[65] S. Bhattacharya, P. Basu, and C. Sajal Bhattacharya, “The southern house mosquito, culex quinquefasciatus: Profile of a smart vector,” J Entomol Zool Stud, vol. 4, no. 2, pp. 73–81, 2016.

[66] D. A. Lapointe, “Dispersal of culex quinquefasciatus (diptera: Culicidae) in a hawaiian rain forest,” Journal of Medical Entomology, vol. 45, no. 4, pp. 600–609, 2008.

[67] WorldPop and Center for International Earth Science Information Network (CIESIN), Columbia University, Global high resolution population denominators project-funded by the bill and melinda gates foundation (opp1134076), 10.5258/SOTON/WP00674, Accessed: [Insert Date], 2018.

[68] S. E. Fick and R. J. Hijmans, “Worldclim 2: New 1km spatial resolution climate surfaces for global land areas,” International Journal of Climatology, vol. 37, no. 12, pp. 4302–4315, 2017.

[69] A. R. North and H. C. J. Godfray, “Modelling the persistence of mosquito vectors of malaria in burkina faso,” Malaria Journal, vol. 17, pp. 1–15, 2018.

[70] Z. Wang and J. Champer, “Optimal spatial release strategies for confined gene drives and wolbachia,” bioRxiv, pp. 2026–03, 2026.

[71] I. K. Kim and P. W. Messer, “Predicting the invasiveness of threshold-dependent gene drives,” bioRxiv, pp. 2025–11, 2025.

[72] D. Florez, R. Cortez, J. M. Hyman, and Z. Qu, “Improving wolbachia-based control programs in urban settings: Insights from spatial modeling,” PLOS Neglected Tropical Diseases, vol. 19, no. 12, e0013787, 2025.

[73] N. H. Barton and M. Turelli, “Spatial waves of advance with bistable dynamics: Cytoplasmic and genetic analogues of allee effects,” The American Naturalist, vol. 178, no. 3, E48–E75, 2011.

[74] X. Zhang, W. Sun, I. K. Kim, P. W. Messer, and J. Champer, “Population dynamics in spatial suppression gene drive models and the effect of resistance, density dependence, and life history,” bioRxiv, 2024.

